# Nocturnin drives mitochondrial NADP(H)/NAD(H) rhythms to regulate steroid rhythm amplitude and time metabolism

**DOI:** 10.64898/2026.07.24.740574

**Authors:** Lauren Palluth, Isara Laothamatas, Tammy-Nhu Nguyen, Emil S. Rasmussen, Lauren G. Zacharias, Mauricio J. Velasquez, Melissa Iñigo-Vollmer, Xiaorong Fu, Thomas P. Mathews, Jeffrey G. McDonald, Shawn C. Burgess, Joseph S. Takahashi, Carla B. Green

## Abstract

Circadian rhythms are conserved biological timekeeping mechanisms crucial for the temporal compartmentalization of metabolic processes. However, the molecular pathways by which circadian rhythms are regulated within metabolism are not fully understood. Nocturnin (NOCT) is a highly rhythmic, clock-controlled NADP(H) phosphatase that has been implicated in numerous metabolic phenotypes. While it is known that NOCT significantly impacts the cellular NADP(H) and NAD(H) pools *in vitro*, NOCT’s impact on their concentrations and rhythmicity *in vivo* has not yet been established. In fact, the rhythmicity of NADH, NADP^+^, and NADPH have yet to be quantified in mammalian nucleated cells. Here, we determined both the whole cell and mitochondrial NAD(H) and NADP(H) rhythms in wild-type and *Noct^-/-^*mouse livers. Unexpectedly, we found a robust rhythm in the mitochondrial NADP(H)/NAD(H) ratio that is antiphase to the respective whole cell rhythm. While loss of NOCT increases the amplitude of the whole cell NADP(H)/NAD(H) rhythm, the mitochondrial rhythm is completely damped in *Noct^-/-^*mice. The constitutively higher relative NADP(H) within *Noct^-/-^*mitochondria drives steroidogenesis, leading to an increased amplitude of plasma corticosterone. Both the acute increase in plasma corticosterone and the disruption of mitochondrial cofactor rhythms caused by loss of NOCT lead to widespread changes in hepatic metabolism. Collectively, we found that NOCT’s control of mitochondrial NADP(H)/NAD(H) rhythms is a novel regulator of steroid amplitude and downstream metabolic rhythms.

## Introduction

Time is an important variable in metabolic health. The time in which an organism eats impacts both its health and lifespan^1–4^. In humans, several clinical trials have shown the timing of treatment has significant impacts on patient outcome^5,6^. Further, the detrimental impact of circadian misalignment by night-shift work, jet lag, and daylight-saving time on metabolic health is well documented^7–11^. Despite this, we still know very little about what regulates metabolic time. Research on what factors impact the phase or amplitude of specific metabolites or pathways can help us not only better understand the drivers of different pathologies but also better design more effective therapeutics.

The core circadian clock is the major timekeeper of metabolism, as the transcription factors CLOCK and BMAL1 directly promote the rhythmic expression of thousands of metabolic genes^12,13^. Notably however, numerous metabolic pathways that are not directly transcriptionally controlled by the core clock still display daily oscillations, suggesting these rhythms are driven by other signaling pathways, such as nutrients from food intake, or are temporally rate-limited by cofactors and products of pathways that are under direct circadian control^14,15^. Nicotinamide adenine dinucleotide (NAD(H)) and its phosphorylated counterpart, nicotinamide adenine dinucleotide phosphate (NADP(H)), are particularly interesting candidates in the study of metabolic time. They are some of the most widely used metabolites in mammalian biology^16^. Further, virtually all the enzymes that regulate the cellular pools of NAD(H) and NADP(H) are under circadian control^17^. This positions NAD(H) and NADP(H) as potential mediators by which the core clock propagates further, indirect control of metabolic rhythms. In fact, previous reports have found daily oscillations of NAD^+^ in murine livers that directly coincide with oscillations in mitochondrial lipid oxidation, protein acetylation, and respiration^18–21^. Despite the importance of NAD^+^ rhythms in the control of metabolic time, no study to date has quantitatively measured how the cellular, or subcellular, pools of NADH, NADP^+^, or NADPH oscillate throughout the day. Further, it is unknown if or how the circadian enzymes that regulate the NAD(H) and NADP(H) pools impact their rhythms.

The clock-controlled NADP(H) phosphatase, nocturnin (NOCT), is of particular interest in addressing these questions^22,23^. NOCT is rhythmically and ubiquitously expressed with peak expression at the onset of the active phase^24–26^. Previous studies have found that *Noct^-/-^* mice have several interesting metabolic phenotypes, including resistance to diet-induced obesity and hepatic steatosis, slowed intestinal chylomicron transit, and disruptions in lipid metabolism pathways^27–29^. In addition, NOCT has two isoforms that differ in both their localization and rhythmicity^30^. The mitochondrial localized isoform of NOCT has a ∼six-fold amplitude rhythm while the cytosolic isoform is constitutively expressed, suggesting that NOCT may differentially regulate distinct subcellular pools of NAD(H) and NADP(H)^31^. Given this, studying NOCT may provide insight into the spatiotemporal regulation of metabolism.

Here, we determined the rhythmicity of the NAD(H) and NADP(H) redox pairs in both the whole cell and mitochondria from wild-type (WT) and *Noct^-/-^* livers. Even though individual dinucleotide species were minimally affected by loss of NOCT, we found a highly robust rhythm in the NADP(H)/NAD(H) ratio within the mitochondria, which is antiphase to the whole cell NADP(H)/NAD(H) rhythm. Strikingly, this mitochondrial rhythm is completely damped in the *Noct^-/-^* mice. The constitutively high relative NADP(H) levels in the *Noct^-/-^* mitochondria drives steroidogenesis, leading to an increased amplitude of plasma corticosterone (CORT). This acute hypercortisolism leads to a temporally discrete upregulation of several CORT-driven pathways, including both gluconeogenesis and lipogenesis. In sum, we found that both the disruption in mitochondrial NADP(H)/NAD(H) oscillations as well as the increased steroid amplitude from loss of NOCT lead to widespread effects on hepatic metabolic rhythms.

## Results

### *Noct^-/-^* mice display altered whole cell and mitochondrial NAD(P)(H) rhythms

The cellular and subcellular rhythmicity of NADH, NADP^+^, and NADPH have yet to be quantified in mammalian nucleated cells. Accurate preservation of redox species requires immediate processing of samples, which is laborious in circadian time series collections. However, with increasing data highlighting the relationship between NAD(P)(H), circadian rhythms, and metabolic health, there is an equally increasing need for quantitative measurements of these cofactors across a circadian cycle. Further, we were particularly interested in determining the contribution of the rhythmic NADP(H) phosphatase, NOCT, to the cofactor rhythms, for loss of NOCT is associated with several metabolic phenotypes^27–29,32^. We hypothesized that NOCT generates cellular NADP(H)/NAD(H) rhythms, and loss of these rhythms leads to metabolic dysfunction.

To measure the oscillations of the NAD(H) and NADP(H) redox pairs quantitatively, we collected livers from both WT and *Noct^-/-^* mice across a circadian cycle (**Figure 1A**). Dinucleotides were immediately extracted from the livers using methods previously optimized to preserve redox state^33^. At the end of the collection, samples were promptly quantified via liquid chromatography-mass spectrometry (LC-MS)^34^. As seen in **Figure 1B**, we found similar NAD^+^ rhythms to what was previously reported in WT mice^19^. While NADH displayed a different profile than its oxidized counterpart, with a singular peak at ZT15, NADP^+^ and NADPH had similar bimodal rhythms, with peaks at both ZT0 and ZT15. Strikingly, NADP(H) oscillated with a nearly two-fold greater amplitude than that of NAD(H). Loss of NOCT resulted in modest effects on the rhythms of the individual species, though significantly damped the ZT15 NADH peak. Even though the pool size of NAD(H) is ∼3-5X larger than the NADP(H) pool, loss of NOCT had a greater effect on lowering cellular NAD(H) than on raising NADP(H) levels. Despite previous reports of daily oscillations in other redox systems, the NAD^+^/NADH and NADP^+^/NADPH redox ratios remained relatively constant throughout the day (**Figure 1B**)^35^. *Noct^-/-^*mice exhibited a more oxidized NAD^+^/NADH ratio throughout the day as compared to WT mice with no effect on the NADP^+^/NADPH ratio. To visualize the enzymatic role of NOCT in the interconversion between NADP(H) and NAD(H), we calculated the NADP(H)/NAD(H) ratio (**Figure 1C**). *Noct^-/-^* mice had a moderately increased relative NADP(H) pool in the active phase, when NOCT is highest in WT mice. Of note, no significant differences in the levels of NADK and NADK2 between genotypes were observed, suggesting that the changes in cofactor levels are in fact due to perturbations in NOCT (**Supplemental Figure 1**). Overall, while we found rhythms in all nicotinamide species in murine liver, NADP(H) displays more robust rhythms than NAD(H). Further, loss of NOCT lowers cellular NAD(H) to a greater extent than it increases NADP(H), suggesting a greater need to maintain specific NADP(H) pools, rhythms, or both than those of NAD(H).

**Figure 1:**
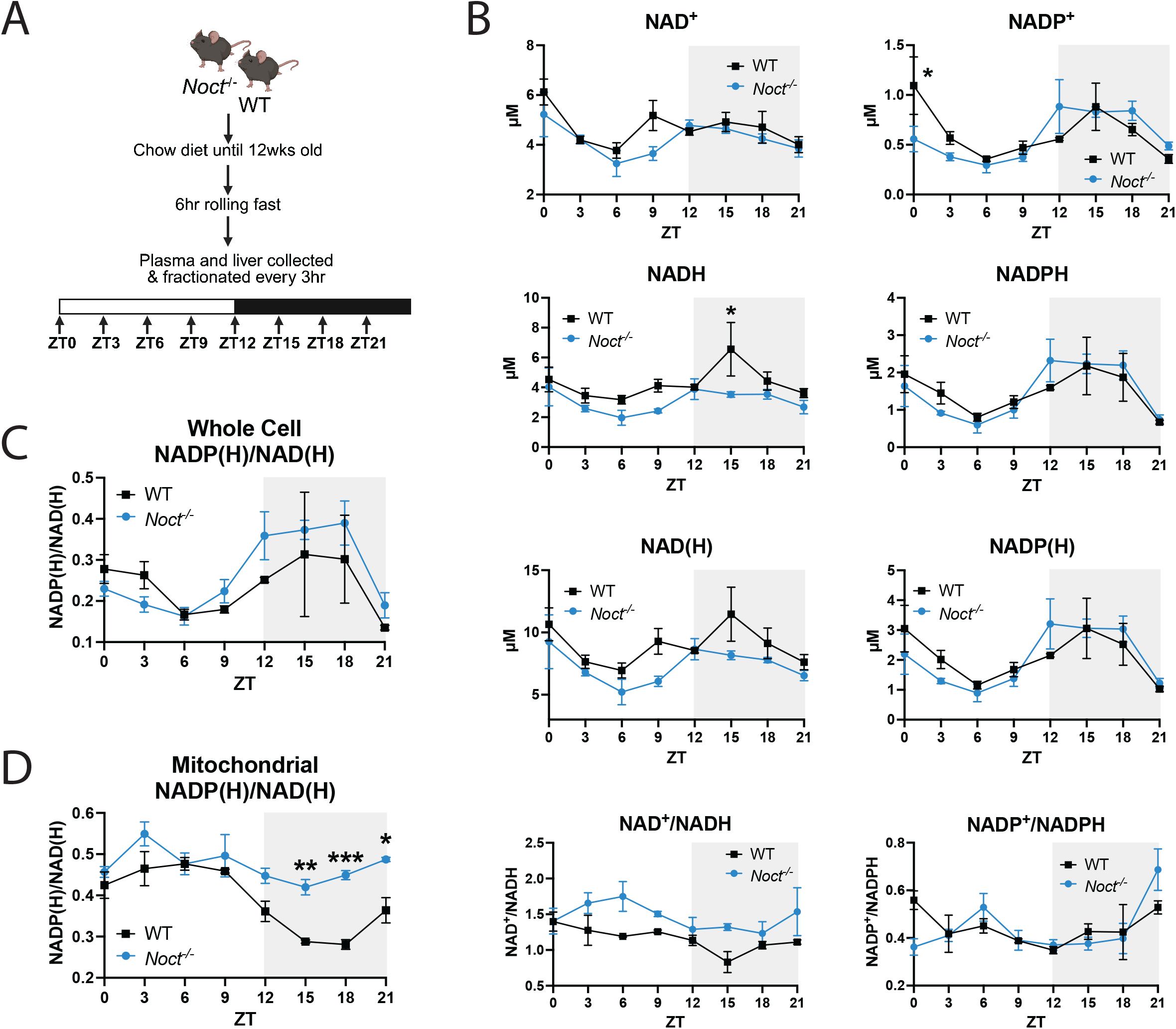
Loss of NOCT impacts cellular and subcellular NAD(P)(H) rhythms. (A) Schematic of circadian tissue collection study design. (B) Nucleotide concentrations in WT and *Noct^-/-^* livers throughout the day (n=3/genotype/timepoint). (C) Ratio of total NADP(H) to NAD(H) in the whole cell. (D) Ratio of total NADP(H) to NAD(H) in the mitochondrial fraction. Two-way ANOVA with multiple comparisons performed on data in (B)-(D). P-value=<0.05 (*), <0.01 (**), <0.001 (***), <0.0001 (****).

Given that the cellular NAD(P)(H) pools are distinctly regulated between organelles and that NOCT’s isoforms differ in both localization and rhythmicity, we hypothesized that subcellular NAD(P)(H) rhythms, and NOCT’s contribution to such, may be masked in our whole cell measurements^31,36–39^. To examine this, we isolated the mitochondrial fraction from WT and *Noct^-/-^*livers and measured NAD(H) and NADP(H). As shown in **Figure 1D**, the mitochondria exhibited a robust NADP(H)/NAD(H) rhythm that peaked in the middle of the rest phase. This is notably antiphasic to the whole cell rhythm, which peaked in the middle of the active phase as shown in **Figure 1C** (**Supplemental Figure 2**). These data suggest that different organelles have distinct NAD(P)(H) rhythms that are masked at the whole cell level. Strikingly, loss of NOCT completely damped the mitochondrial rhythm. Compared to WT, *Noct^-/-^*mitochondria maintained significantly higher NADP(H) levels relative to NAD(H) throughout the active phase. Collectively, our data suggest that NOCT drives mitochondrial NADP(H)/NAD(H) rhythms, and loss of NOCT leads to significantly less conversion of NADP(H) to NAD(H) in the active phase.

### NOCT regulates the circadian amplitude of steroid hormones

To assess the physiological relevance of the higher relative NADP(H) in the *Noct^-/-^* mitochondria, we determined whether NADP(H)-dependent pathways were upregulated in the *Noct^-/-^* mice. The majority of steroid hormones display robust circadian and ultradian rhythms, and glucocorticoids can entrain peripheral clocks ^40–46^. Thus, we hypothesized that NOCT-mediated regulation of mitochondrial NADP(H)/NAD(H) rhythms regulates steroid rhythms (**Figure 2A**), creating a cascade to downstream metabolic rhythms.

**Figure 2:**
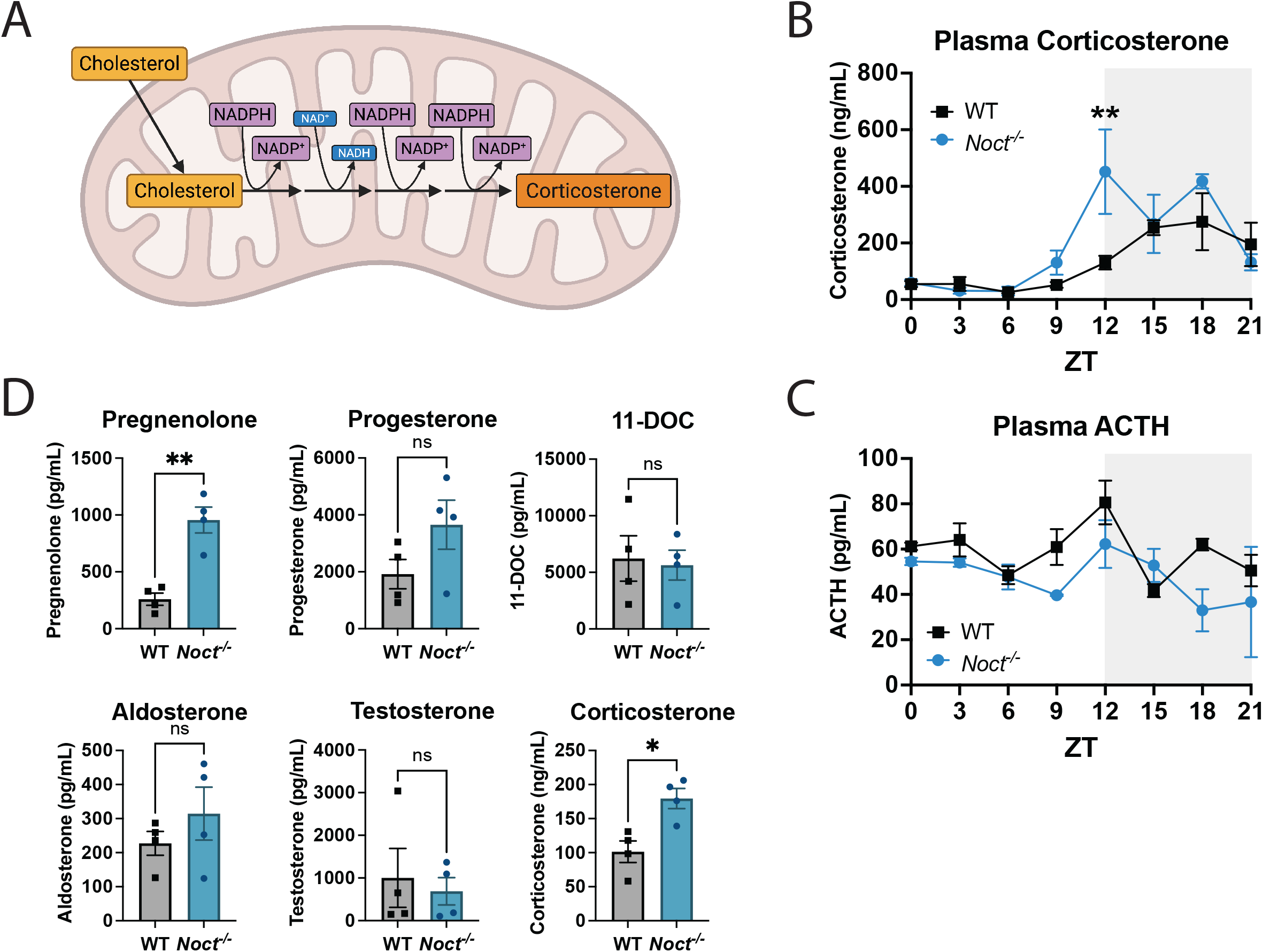
Loss of NOCT increases plasma steroid rhythm amplitude. (A) Schematic of CORT synthesis from cholesterol, highlighting the main cofactors used in each reaction. (B) Plasma CORT in WT and *Noct^-/-^* mice throughout the day (n=3/genotype/timepoint). Two-way ANOVA with multiple comparisons performed. (C) Plasma ACTH in WT and *Noct^-/-^* mice throughout the day (n=3/genotype/timepoint). Two-way ANOVA with multiple comparisons performed. (D) Panel of plasma steroids in WT and *Noct^-/-^*mice (n=4/genotype). Welch’s T-test performed. P-value=<0.05 (*), <0.01 (**), <0.001 (***), <0.0001 (****).

To determine the effect of NOCT on steroidogenesis, we first measured CORT from the *Noct^-/-^* and WT circadian plasma samples. CORT is a glucocorticoid with widespread effects on metabolism and exhibits a robust circadian rhythm in phase with NOCT, as the levels of both CORT and NOCT peak at the onset of the active phase^24,45,47,48^. Remarkably, we found that *Noct^-/-^* mice have a significantly higher amplitude of plasma CORT than WT mice (**Figure 2B**). As the plasma half-life of CORT is ∼66 minutes, we postulated that this acute increase in circulating CORT was either due to increased upstream hypothalamic-pituitary-adrenal (HPA) axis signaling or increased synthesis due to high mitochondrial NADP(H)^49^. To test if HPA axis signaling is perturbed in the *Noct^-/-^*mice, we measured the concentration of adrenocorticotropic hormone (ACTH) in the circadian plasma samples. As seen in **Figure 2C**, *Noct^-/-^* mice exhibited similar plasma ACTH levels to WT throughout the day, suggesting increased upstream signaling was not driving the higher CORT amplitude. To test whether steroid synthesis is impacted by loss of NOCT, we measured a panel of steroids from *Noct^-/-^* and WT plasma collected at ZT14 in order to capture the circadian peak of multiple steroids. As seen in **Figure 2D**, we found significantly higher steroidogenesis precursors in the *Noct^-/-^* plasma. Plasma pregnenolone was markedly greater in *Noct^-/-^* than WT mice, consistent with the increased NADP(H) availability supporting the mitochondrial conversion of cholesterol to pregnenolone, the rate-limiting step in steroidogenesis^50,51^. While we also found a moderate increase in aldosterone, a hormone synthesized from CORT, we did not observe any differences in testosterone, possibly suggesting that loss of NOCT has a greater effect on steroids synthesized primarily within the mitochondria^51,52^. As previously mentioned, glucocorticoids can entrain peripheral clocks^46^. However, there were no remarkable differences in the phase nor levels of core clock proteins in WT and *Noct^-/-^* livers (**Supplemental Figure 1B**). In summary, we have found that loss of NOCT increases the amplitude of plasma CORT. This acute rise in CORT is likely due to increased steroidogenesis, driven by increased NADPH availability within the mitochondria.

### Loss of NOCT impacts glucose homeostasis in a time-of-day-dependent manner

NOCT’s control of plasma steroid amplitude potentially expands its influence on metabolic rhythms beyond the mitochondria. Specifically, CORT is known to have pleiotropic effects on metabolism^53^. Pathologies of hypercortisolism, such as Cushing’s Disease, lead to obesity, increased lipolysis, hyperglycemia, and insulin resistance^54,55^. However, *Noct^-/-^* mice are resistance to excessive weight gain and remain lean on high-fat diets (HFD), suggesting the temporally distinct increase in plasma CORT elicits a pathologically different response than the constitutively high CORT seen in Cushing’s ^27,28^. Given this, we were interested to see if *Noct^-/-^* mice display other phenotypes associated with hypercortisolism, such as disruptions in glucose and lipid metabolism.

As outlined in **Figure 3A**, we first measured fasting glucose in male and female WT and *Noct^-/-^*mice at both ZT14, during the peak of CORT and NOCT, and at ZT2, during their trough. Strikingly, while there were no differences in blood glucose between genotypes at ZT2, both male and female *Noct^-/-^* mice exhibited significantly higher fasting glucose at ZT14 than WT mice (**Figure 3B**). To test whether this hyperglycemia correlates with increased CORT, we challenged the mice with short-term HFD, as this acute stress is known to increase plasma CORT^56^. As seen in **Figure 3C**, the difference in fasting glucose at ZT14 was exacerbated after both 1- and 3-weeks HFD. Accordingly, this exacerbation mirrored the difference we see in plasma CORT between genotypes (**Figure 3D**).

**Figure 3:**
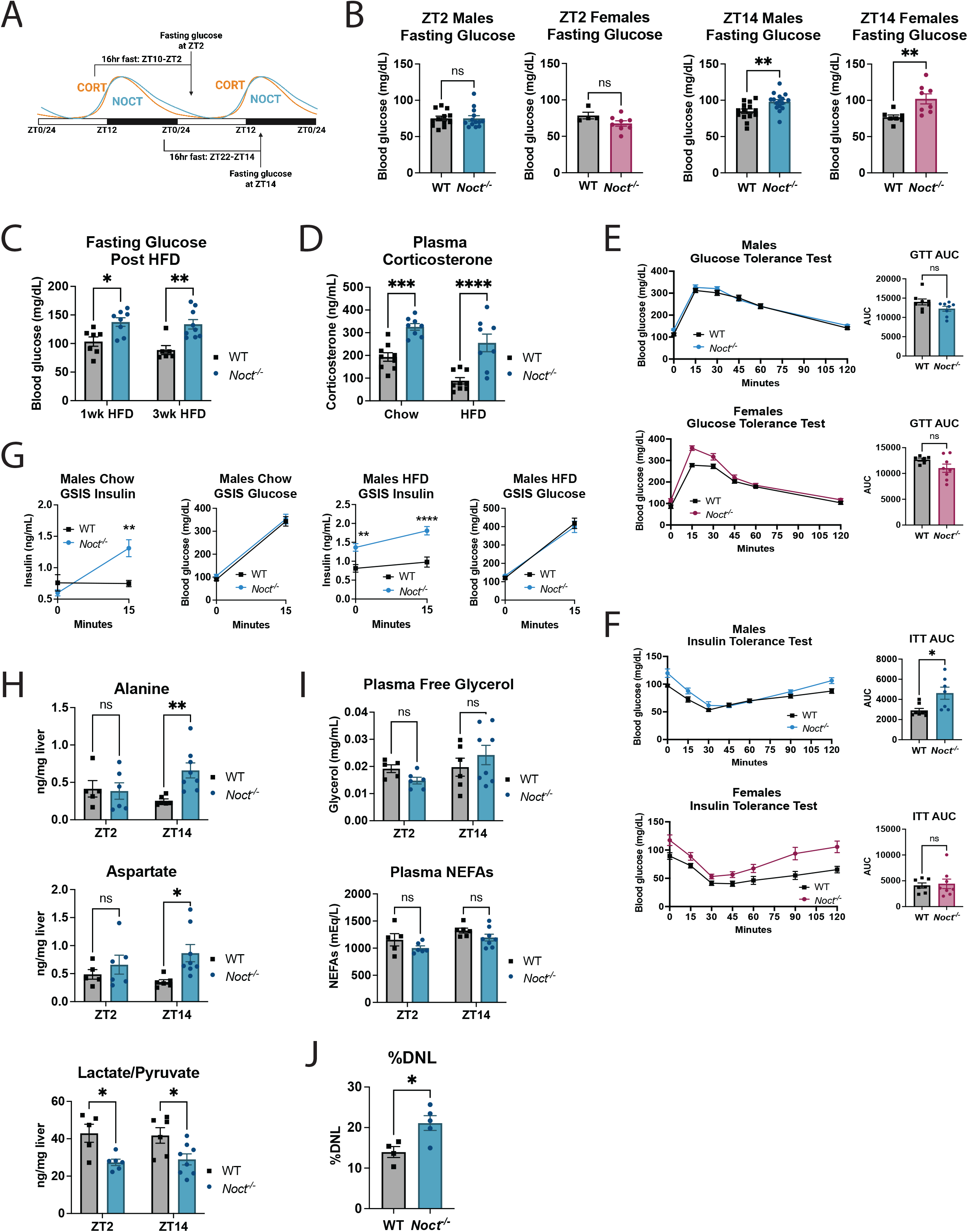
Loss of NOCT perturbs glucose and lipid metabolism in a time-of-day dependent manner. (A) Experimental overview of fasting glucose measurements. Both cohorts were fasted for 16-hours directly prior to blood collection from either ZT10-ZT2 or ZT22-ZT14. (B) WT and *Noct^-/-^* mice were fed *ad libitum* chow diet, fasted 16hr, then blood glucose measurements were taken at either ZT2 or ZT14. For the ZT2 cohort, male WT n=12 and *Noct^-/-^* n=13; female WT n=4 and *Noct^-/-^* n=8. For the ZT14 cohort, male WT n=14 and *Noct^-/-^* n=16; female WT n=7 and *Noct^-/-^* n=8. Welch’s t-test performed. (C) Fasting glucose measurements from WT and *Noct^-/-^* mice at ZT14 after either 1- or 3-weeks HFD followed by 16-hour fast (n=6 for WT; n=8 for *Noct^-/-^*). Two-way ANOVA with multiple comparisons performed. (D) Plasma CORT measurements from WT and *Noct^-/-^* mice at ZT14 after either chow or 3-weeks HFD followed by 16-hour fast (n=9 for WT; n=8 for *Noct^-/-^*). Mixed-effect analysis with multiple comparisons performed. (E) IPGTT of male and female WT and *Noct^-/-^* mice at ZT14 post 16-hour fast (n=8 for WT and *Noct^-/-^* males; n=7 for WT and n=8 *Noct^-/-^* females). Welch’s t-test performed on the AUC. (F) IPITT of male and female WT and *Noct^-/-^* mice at ZT14 post 16-hour fast (n=8 for WT and n=7 *Noct^-/-^* males; n=7 for WT and n=8 *Noct^-/-^* females). Welch’s t-test performed on the AUC. (G) Plasma insulin and blood glucose measurements during GSIS test on WT and *Noct^-/-^* male mice at ZT14 after either chow or 3-weeks HFD followed by a 16-hour fast (n=7 WT and n=8 *Noct^-/-^*). (H) Hepatic alanine and aspartate concentrations WT and *Noct^-/-^* mice collected at either ZT2 or ZT14. Mice were fed *ad libitum* facility chow diet then fasted for 16-hours prior to each timepoint. For the ZT2 cohort, WT n=5 and *Noct^-/-^* n=6. For the ZT14 cohort, WT n=6 and *Noct^-/-^* n=8. Two-way ANOVA with multiple comparisons performed. (I) Plasma free glycerol and NEFAs measured from WT and *Noct^-/-^* mice described in (H). Two-way ANOVA with multiple comparisons performed. (J) Fractional DNL/2hr in n=4 WT and n=5 *Noct^-/-^* male livers. Mice were injected with ^2^H_2_O at ZT12, provided with 6% ^2^H_2_O drinking water, and livers were collected at ZT14. Welch’s t-test performed. P-value=<0.05 (*), <0.01 (**), <0.001 (***), <0.0001 (****).

To assess if *Noct^-/-^* mice display larger perturbations in glucose homeostasis, we performed a series of tolerance tests. Interestingly, though CORT has been shown to decrease glucose uptake, *Noct^-/-^*mice displayed similar glucose clearance to WT mice (**Figure 3E**). However, *Noct^-/-^* male mice exhibited moderate insulin resistance, as seen in **Figure 3F**, yet *Noct^-/-^* female mice did not, suggesting a sex-specific response. Despite apparent insulin resistance, *Noct^-/-^* male mice retained beta-cell function with marked glucose-stimulated insulin secretion at ZT14 under both chow and short-term HFD conditions (**Figure 3G**). As the fasting hyperglycemia could not be explained by defects in glucose uptake or beta-cell function, we then investigated if gluconeogenesis was upregulated in the *Noct^-/-^*mice. We collected livers from *Noct^-/-^* and WT mice at both ZT2 and ZT14 and measured a panel of gluconeogenic organic and amino acids^57^. As seen in **Figure 3H**, *Noct^-/-^* mice had significantly higher levels of hepatic alanine and aspartate at ZT14, when we observed the hyperglycemia, but not at ZT2, when we observed no differences in blood glucose. Further, *Noct^-/-^* mice exhibited significantly lower lactate/pyruvate ratio, which is a measure of unbound NADH/NAD^+^, at both timepoints (**Figure 3H**). Of note, this measure parallels data in **Figure 1B** where the *Noct^-/-^*mice had a more oxidized NADH/NAD^+^ ratio from measures of the total NAD(H) pool (bound and unbound). Previous studies have found that lower hepatic NADH/NAD^+^ drives gluconeogenesis^58–60^. Thus, the lower lactate/pyruvate ratio, along with the time-dependent increase in substrate availability, observed in *Noct^-/-^*mice support the notion of increased gluconeogenesis. In summary, we have shown that loss of NOCT leads to fasting hyperglycemia in a time-of-day-dependent manner. The fasting hyperglycemia is likely due to a temporally distinct increase in gluconeogenesis that is driven by increased CORT amplitude.

Glucocorticoids have also been shown to increase both lipolysis and lipogenesis ^48,55,61–64^. To first gauge the extent of lipolysis, we collected plasma from *Noct^-/-^*and WT mice at both ZT2 and ZT14. As seen in **Figure 3I**, there were no remarkable differences in plasma free glycerol nor non-esterified fatty acids (NEFAs), suggesting no or minimal effect on basal lipolysis. As loss of NOCT did not significantly increase whole cell NADP(H), we then determined if *de novo* lipogenesis (DNL) was altered in *Noct^-/-^*mice. To measure lipogenic flux, we administered deuterated water (^2^H_2_O) to *Noct^-/-^* and WT mice at ZT12 and subsequently collected their livers at ZT14. Using gas chromatography high resolution orbitrap mass spectrometry (GC-HRMS) to quantify ^2^H incorporation in palmitate, we found significantly higher DNL in *Noct^-/-^* livers (**Figure 3J**)^65,66^. These data demonstrate that loss of NOCT has no impact on lipolysis yet significantly increases hepatic DNL. Overall, NOCT’s regulation of steroid amplitude propagates its influence on metabolic rhythms beyond the mitochondria to both hepatic glucose and lipid metabolism.

### Mitochondrial NOCT is sufficient to reverse the hyperglycemia and hypercortisolism observed in *Noct^-/-^* mice

To test whether the phenotypes we observed were in fact due to increased mitochondrial NADP(H)/NAD(H), we generated a CRISPR knock-in mouse line in which the downstream initiating methionine of *Noct* was mutated to alanine (M65A), forcing all translation of *Noct* to include the mitochondrial targeting sequence (**Figure 4A**)^30,31^. This novel *Noct^mito^* mouse line does not express cytosolic NOCT but does express the mitochondrial NOCT isoform. To confirm the mitochondrial NADP(H)/NAD(H) was not increased in these mice, we isolated the mitochondrial fraction from livers at ZT14 and measured the dinucleotides from the mitochondrial pellet. As seen in **Figure 4B**, *Noct^mito^* mice did not exhibit high relative NADP(H), demonstrating that mitochondrial NOCT was sufficient to rescue the rhythms in mitochondrial NADP(H). We also confirmed that the *Noct^mito^* mice had comparable levels of NOCT in their mitochondria to WT mice (**Supplemental Figure 3A**).

**Figure 4:**
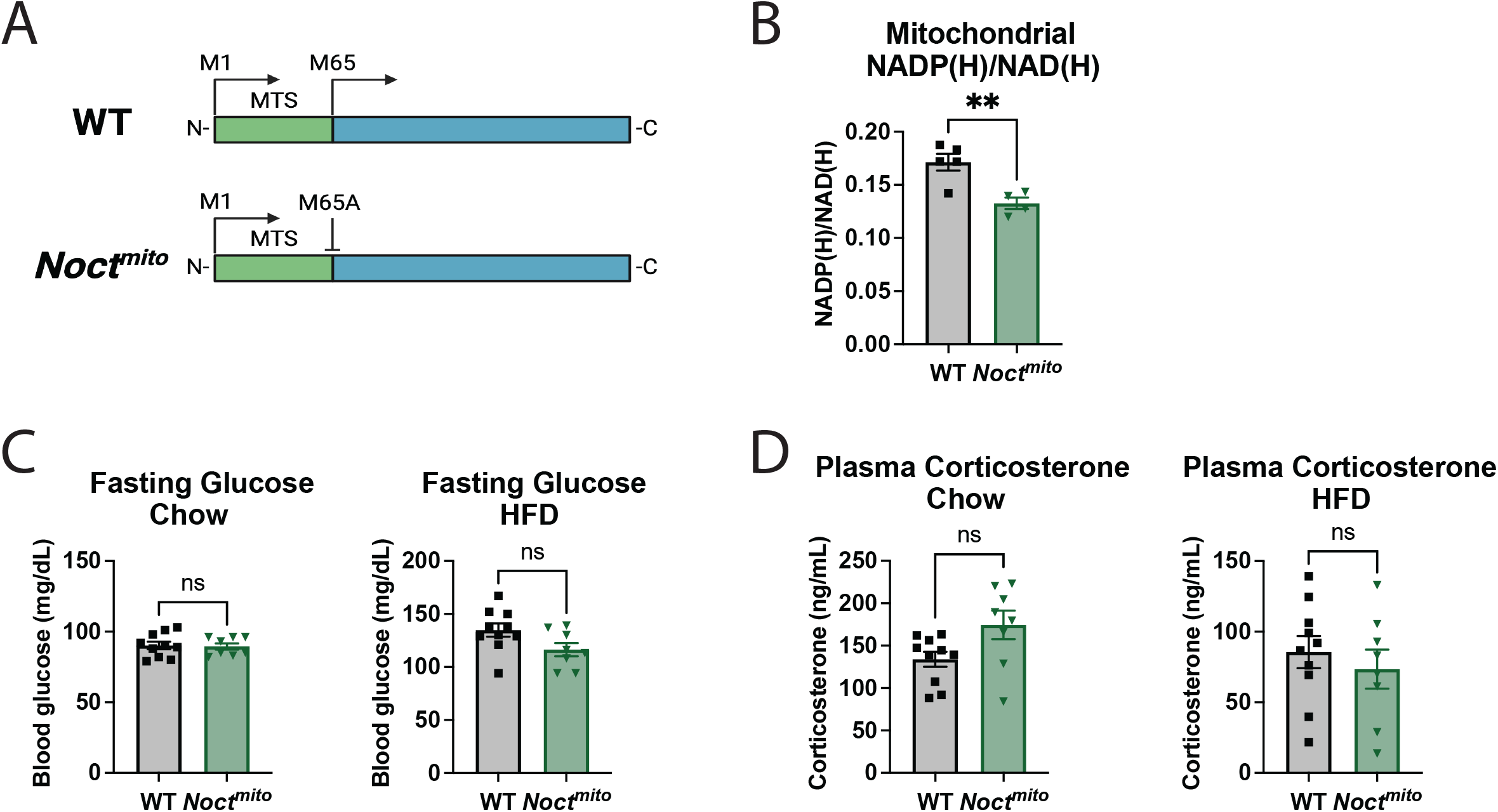
Restoration of mitochondrial NADP(H)/NAD(H) reverses the hypercortisolism and hyperglycemia seen in the *Noct^-/-^* mice. (A) Schematic of the WT *Noct* sequence to highlight the two in-frame methionines. Methionine (M65) is mutated to alanine in the *Noct^mito^*mouse line. (B) Ratio of NADP(H) to NAD(H) in the isolated mitochondrial fraction of WT and *Noct^mito^* male livers collected at ZT14 post 16-hour fast (n=5 WT and n=4 *Noct^mito^*). Welch’s t-test performed. (C) Fasting glucose at ZT14 from WT and *Noct^mito^*males. Mice were *ad libitum* fed chow or 3-weeks HFD then fasted for 16-hours prior to measurements (n=10 WT and n=8 *Noct^mito^*). Welch’s t-test performed. (D) Plasma CORT at ZT14 from WT and *Noct^mito^* males described in (C). Welch’s t-test performed. P-value=<0.05 (*), <0.01 (**), <0.001 (***), <0.0001 (****).

Using this genetic tool, we investigated if the hypercortisolism and hyperglycemia we observed in *Noct^-/-^* mice were reversed with rescue of NOCT only in the mitochondria and the resulting lower relative mitochondrial NADP(H). We measured plasma CORT in *Noct^mito^* and WT mice at ZT14 following both chow and 3-week HFD exposure. As seen in **Figure 4C**, *Noct^mito^* mice did not have high plasma CORT in either condition. We then sought to determine whether *Noct^mito^* mice would display hyperglycemia in the absence of high mitochondrial NADP(H) or high CORT. As predicted, the *Noct^mito^* mice did not exhibit high blood glucose at ZT14 on either chow or short-term HFD (**Figure 4D**). Further, *Noct^mito^* mice maintained normal glucose clearance and insulin sensitivity (**Supplemental Figure 4A-B**). Our data suggest that restoration of NOCT in the mitochondria, and thus restoration of mitochondrial NADP(H)/NAD(H), is sufficient to reverse the hypercortisolism, hyperglycemia, and insulin resistance observed in the global *Noct^-/-^* mice.

### Loss of NOCT widely disrupts metabolic rhythms

Having shown that loss of NOCT impacts both mitochondrial and non-mitochondrial pathways, we sought to gain a broader understanding of NOCT’s impact on metabolic rhythms. Therefore, we performed untargeted metabolomics from *Noct^-/-^* and WT livers collected across a circadian cycle (**Figure 1A**). Metabolites were immediately extracted from the livers at each timepoint and subsequently quantified via LC-MS. To visualize how loss of NOCT impacted metabolomic rhythmicity, we performed an unbiased correlation network analysis using the principles of Weighted Gene Co-expression Network Analysis (WGCNA)^67^. This analysis groups the metabolites based on their abundance pattern into five distinct modules assigned different colors (**Figure 5A**). **Figure 5B** highlights a subset of metabolites significantly altered in *Noct^-/-^* mice. The blue-colored metabolites represent the “Blue” module, which had decreased levels in *Noct^-/-^*livers in the active phase, the yellow-colored metabolites represent the “Yellow” module, which had increased levels in *Noct^-/-^* livers in the active phase, so on and so forth (**Figure 5A-B**). Interestingly, metabolites in the same pathway largely grouped within the same module, suggesting that pathways are uniquely temporally affected by changes in NAD(H) and NADP(H) rhythms.

**Figure 5:**
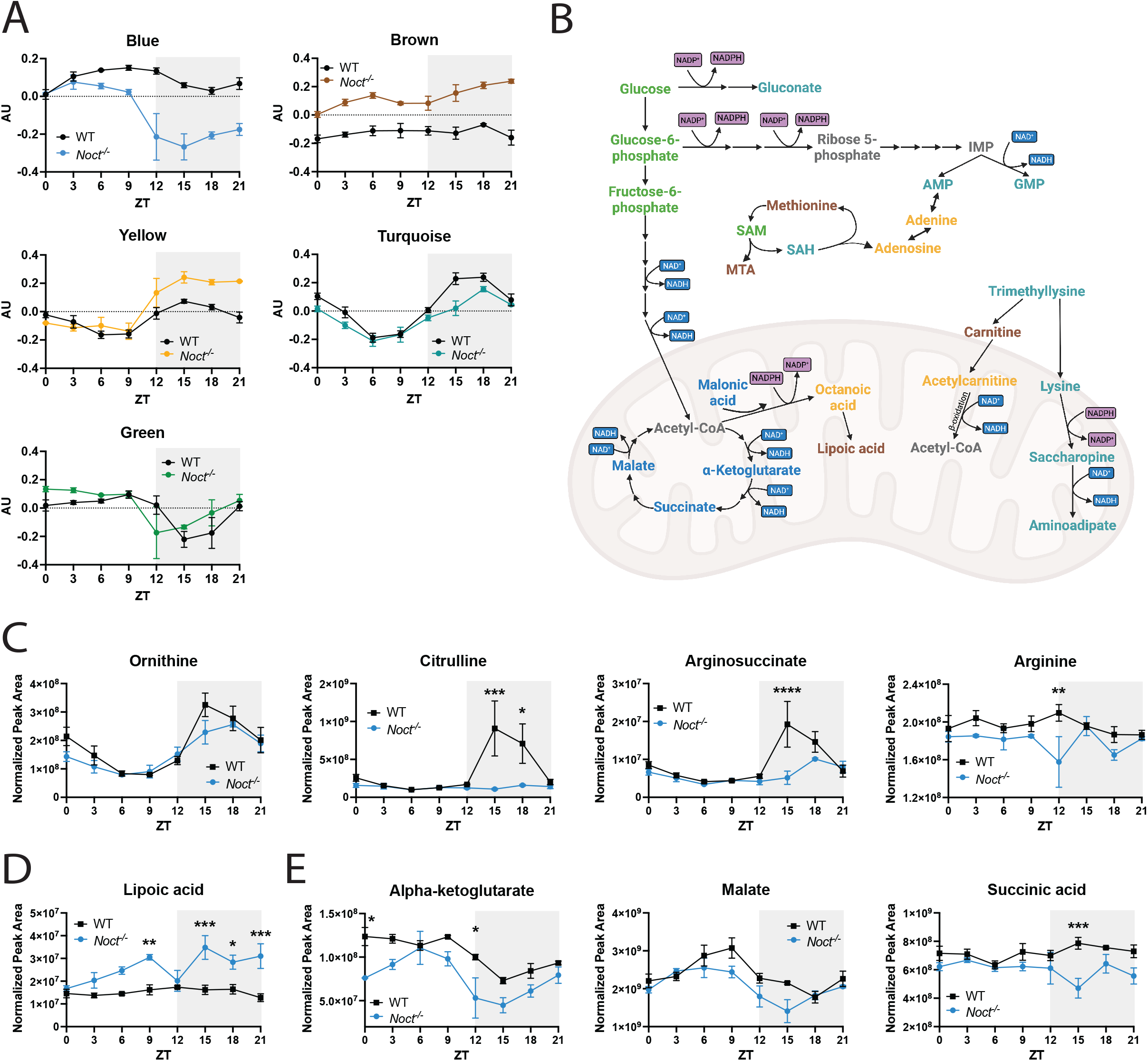
Loss of NOCT impacts hepatic metabolic rhythms. (A) Eigenvalues representing modules generated by the grouping analysis. (B) Schematic of metabolic pathways highlighting subsets of metabolites that grouped into the modules represented in (A). (C) Select urea cycle metabolites. (D) Select TCA cycle metabolites. (E) Lipoic acid. Two-way ANOVA with multiple comparisons performed on graphs (C)-(E). P-value=<0.05 (*), <0.01 (**), <0.001 (***), <0.0001 (****).

As expected, these data suggest an overall downregulation of mitochondrial NAD(H)-dependent processes and upregulation of mitochondrial NADP(H)-dependent processes. Specifically, metabolites in the TCA cycle grouped in the “Blue” module and largely displayed decreases and phase shifts in *Noct^-/-^* livers (**Figure 5A-B,E**). We also observed a dramatic damping of hepatic citrulline and downstream urea cycle metabolites, arginosuccinate and arginine, yet unremarkable effects on upstream ornithine (**Figure 5C**). The conversion of ornithine to citrulline is an ATP-dependent, mitochondrial reaction, and low hepatic citrulline is a marker of reduced complex I activity^68^. However, products of NADP(H)-dependent pathways, such as octanoic acid and lipoic acid of the mitochondrial fatty acid synthesis (mtFAS) pathway, grouped in the “Yellow” and “Brown” modules and were significantly increased in *Noct^-/-^* livers (**Figure 5A-B,D**). Overall, these data support the notion that loss of NOCT leads to a reduction in products from mitochondrial NAD(H)-dependent pathways and an increase in products from NADP(H)-dependent pathways.

Interestingly, loss of NOCT also altered the rhythms of numerous metabolites nonexclusive to the mitochondria. We saw pronounced damping of all monophosphate rhythms, as the amplitude of uridine, cytidine, guanosine, inosine, and adenine monophosphate were all significantly lower in *Noct^-/-^* livers (**Supplement Figure 5A**). However, we observed an accumulation of the respective nucleosides and nucleotides in *Noct^-/-^* livers (**Supplemental Figure 5B**). There were also notable differences in one-carbon metabolism, with time-of-day-dependent decreases in S-adenosylmethionine and S-adenosylhomocysteine as well as a significant increase in 5’-S-methyl-5’-thioadenosine throughout the day (**Supplemental Figure 5C**). As these metabolites are neither exclusively mitochondrial nor CORT-regulated, these data highlight how the change in mitochondrial NADP(H)/NAD(H) rhythms due to loss of NOCT has the capacity to induce widespread effects on metabolic timing.

## Discussion

NAD(H) and NADP(H) are some of the most ubiquitously used cofactors in mammalian biology^16^. The circadian control of all the major enzymes regulating the cellular NAD(P)(H) pools raised questions regarding the rhythmicity of these cofactors as well as their potential role in propagating the temporal cues of the core clock to peripheral metabolism. Further, the compartment-specific regulation of NAD(P)(H) pools suggested that the rhythmicity of the cofactors may vary between organelles. The findings presented in our study confirm the rhythmicity of NAD(P)(H), how its rhythmicity is distinct within the mitochondria, and how this drives downstream metabolic rhythms.

While some enzymes can utilize either NAD(H) or NADP(H), these cofactors are largely involved in distinct pathways. NAD(H) is utilized in most catabolic reactions as well as in deacetylation and ADP-ribosylation of proteins and nucleic acids; however, NADP(H) is primarily utilized in anabolic reactions and antioxidant defense^17^. It, therefore, makes evolutionary sense that the availability of these cofactors would be spatially and temporally discrete. As we found in WT mitochondria, NAD(H) is relatively higher during the active phase. This likely contributes to the TCA cycle and beta-oxidation, as the flux through these pathways are higher in the active, fed state. The drop in NAD(H) in relation to NADP(H) during the rest phase likely switches mitochondrial metabolism to a state more focused on oxidant defense and repair. NOCT has a significantly more robust rhythm within the mitochondria than the cytosol, consistent with the notion that the mitochondria are the major metabolic signaling hub^69,70^. Our data suggest that the robust mitochondrial rhythm of NOCT creates a temporal division between NAD(H) and NADP(H) availability, which increases the energetic efficiency of mitochondrial metabolism. This finding presents a novel perspective of how circadian rhythms within metabolism are generated.

Importantly, we found an antiphase relationship between the mitochondrial NADP(H)/NAD(H) rhythm, which peaks in the middle of the rest phase, and the whole cell NADP(H)/NAD(H) rhythm, which peaks in the middle of the active phase. These findings are particularly remarkable given that the whole cell lysate contains the mitochondrial fraction that is likely diluting the antiphasic pattern. The mitochondrial NAD(P)(H) pool is dependent on NAD^+^ import via the SLC25A51 transporter^36,71,72^. However, the total mitochondrial NAD(P)(H) pool did not differ between genotypes throughout the day, implying there are no defects nor compensatory effects on NAD^+^ import in *Noct^-/-^* mice. Though the subcellular pools are discrete, recent data has highlighted their interconnectivity with the mitochondrial pool as the major buffer^73^. Thus, further studies are required to determine the effect of NOCT on other subcellular NAD(P)(H) pools and rhythms.

Our whole cell measurements revealed that loss of NOCT lowers cellular NAD(H) to a greater extent than it raises cellular NADP(H) (**Figure 1B**). This finding is thought provoking, for logic would assume that loss of the phosphatase would have a larger impact on the smaller cofactor pool. A compensatory decrease in NADK activity could partially explain this effect. Consistent with this possibility, product inhibition of NADK by NADPH has been shown *in vitro*^74^. Moreover, because NADK preferentially phosphorylates NAD^+^ over NADH, reduced NADK activity would theoretically lead to a higher NAD^+^/NADH ratio, which we do in fact see in *Noct^-/-^* livers^75^. However, previous studies reported no significant differences in the NAD^+^/NADH ratio upon NADK deletion, and NADK levels were similar in *Noct^-/-^* and WT livers in this study (**Supplemental Figure 1B**)^75,76^. An alternative explanation may be a reduction in NAD^+^ synthesis due to loss of NOCT; however, the total NAD(P)(H) pool did not substantially differ between genotypes throughout the day. Since our data has shown that NAD(P)(H) rhythms differ between the whole cell and mitochondria, future studies measuring the cofactor rhythms in other organelles will be important for defining the differential impact of NOCT on NAD(H) versus NADP(H). Studies examining the compensatory relationship between NADK, NADK2, and NOCT as well as their impact on regulating NAD^+^ synthesis would also aid our understanding of these data.

Much attention has focused on how the cofactor’s redox state affects the directionality of pathways. While there is ample data supporting this claim, we did not detect any rhythms in the NAD^+^/NADH nor NADP^+^/NADPH ratios^58,77–79^ (**Figure 1B**). This finding is especially interesting given that the reduced and oxidized species differentially regulate CLOCK:BMAL1 and NPAS2:BMAL1 binding to DNA^80^. Since redox rhythms likely differ between tissues, quantifying these rhythms in other tissues, such as the SCN, is necessary. Additionally, our extraction method dissociates the bound cofactors, raising the possibility that a redox rhythm would only be observed when measuring unbound pools. We did not observe any variation between the ZT2 and ZT14 timepoints in the lactate/pyruvate ratio, a surrogate measure of the unbound NADH/NAD^+^ ratio; however, sampling only two timepoints is insufficient to exclude rhythmicity (**Figure 3H**). The cofactor redox state may govern more transient changes in metabolism while phosphorylation state may govern larger metabolic shifts. Moreover, if redox rhythms are driven solely by food intake, the rolling fast used in our study design may have masked the rhythms. Future studies measuring NAD(P)(H) rhythms in a time-restricted feeding paradigm as well as quantification of unbound pools are needed to address these questions^81^.

Our results also reveal a novel mechanism regulating plasma steroid amplitude. To our knowledge, no other genetic models have been shown to temporally and acutely increase steroidogenesis. Therefore, NOCT is now the only known genetic regulator of steroid amplitude. From an evolutionary perspective, it is peculiar that a NADP(H) phosphatase would be in-phase with the NADP(H)-dependent reactions of steroidogenesis. However, this suggests a requirement to moderate the rise in plasma steroids, providing another possible mechanism to safeguard the energetic efficiency of metabolism. Interestingly, high CORT amplitude has been observed in models of caloric-restriction^82^. Further, studies have found that high amplitude CORT, induced by timed CORT injections, does not elicit the same deleterious effects on physiology seen with constitutively high CORT^83–86^. Thus, the rhythmicity of steroids is more important factor for metabolic health than their overall levels. Given the increasing evidence showing the negative impacts of chronically high glucocorticoids in both health and lifespan, studies examining how loss of NOCT affects lifespan would provide valuable insight on how both glucocorticoids and NAD(P)(H) impact longevity^87^.

Loss of NOCT markedly altered hepatic metabolite rhythms. The disruption in metabolic timing is likely due to a combined effect from altered cofactor rhythms as well as increased CORT amplitude. Lower TCA cycle metabolites observed in *Noct^-/-^* livers could be a result of lower mitochondrial NAD^+^ or increased demands for gluconeogenic and lipogenic precursors (**Figure 5E**). The significant damping of monophosphate rhythms along with increased levels of respective nucleosides and nucleotides highlights disruption in purine and pyrimidine metabolism (**Supplemental Figure 5A-B**). We additionally found striking differences in the rhythms of folate cycle metabolites (**Supplemental Figure 5C**). While these data suggest larger changes in one-carbon metabolism, future studies utilizing metabolic tracers are necessary to understand how the rhythmic flux through these pathways are impacted by NOCT.

In summary, we have found that NOCT drives mitochondrial NADP(H)/NAD(H) rhythms. Loss of NOCT leads to higher relative mitochondrial NADP(H) in the active phase, which acutely increases steroidogenesis and the amplitude of plasma CORT. The higher amplitude CORT leads to an upregulation of some, but not all, pathways known to be increased in pathologies of hypercortisolism. Additionally, loss of NOCT perturbs mitochondrial timing beyond steroidogenesis, as we found an overall decrease in products of NAD(H)-dependent pathways and increase in products from NADP(H)-dependent pathways. As NOCT is clock-controlled, these data offer a new perspective on how circadian rhythms are generated in peripheral metabolism.

## Methods

### Circadian collection: husbandry, tissue and plasma collection, NAD(P)(H) and metabolite extraction

N=24/genotype WT and *Noct^-/-^* male mice were single-housed starting at 8-weeks of age and maintained in 12:12 light:dark (LD) conditions and facility chow diet until they were 12-weeks old. For the collection, all mice were kept in LD conditions such that any observed rhythms reflect physiological fluctuations. Additionally, each set of mice were fasted (removal of food and cage change) 6-hours prior to their respective timepoint in order to minimize variability due to food intake while still minimizing stress caused by a long-term fast. N=3/genotype were sacrificed every 3-hours. To preserve redox state, mice were sacrificed by live decapitation followed by immediate trunk blood collection into EDTA-coated tubes (Fisher NC9990563) and excision of the liver. Blood samples were later spun at 2,000xg for 15-minutes at 4°C. Resulting plasma was aliquoted into new tubes and stored at -80°C until later analysis. Approximately 100 mg chunks of liver were promptly put in tubes containing one stainless steel bead (Qiagen 69989) and 1 mL of 40:40:20 acetonitrile:methanol:water (ACN Fisher Scientific A955-4; MeOH Fisher Scientific A456-4) with 0.1 M formic acid (Fisher Scientific A117-50) for NAD(P)(H) measurements or 1 mL of 80% methanol for untargeted metabolomics and then immediately homogenized with a MP Biomedicals FastPrep-24 5G lysis system. Samples used for NAD(P)(H) measurements were then neutralized with NH_4_HCO_3_ as described previously^33^. All samples were then transferred to a 1.5 mL tube and centrifuged at high speed (∼20,160xg) for 15-minutes at 4°C. Supernatant was transferred to a new tube and protein concentration was quantified with Pierce BCA assay (Thermo Scientific #23225). Metabolomics samples were diluted to 0.5 mg/mL in 80% MeOH, and NAD(P)(H) samples were diluted to 0.1 mg/mL in the 40:40:20 mixture. Both sets were then vortexed and spun again at high speed for 15-minutes at 4°C. Supernatant was transferred to LC-MS vials and 10 μL of 20 μM internal standards were added to the NAD(P)(H) samples.

All metabolites were analyzed by LC-MS on an Orbitrap Exploris 480 mass spectrometer coupled to a Vanquish Flex liquid chromatography system (Thermo Scientific, Bremen, DE) equipped with a ZIC-pHILIC column (Millipore-Sigma, St Louis, MO). All data were acquired as previously described^88,89^. For global metabolomics, data from individual samples were acquired with a polarity-switching full scan acquisition method. Pooled samples were created from each individual condition and acquired with a data-dependent MS/MS acquisition method. Data files were processed with Compound Discoverer 3.5 (Thermo Scientific) and searched against in-house spectral databases, metabolite libraries made from purified standards, and mzCloud, an online spectral database maintained by Thermo Scientific. All metabolites were carefully reviewed by analytical chemists to ensure accurate identification and integration. NADP^+^ and NADPH were acquired with a targeted selected ion monitoring method as described previously^34^. Peaks were integrated and quantified using TraceFinder 5.2 (Thermo Scientific).

### Fractionation

Fractionation methods were previously described^31^. Briefly, livers were homogenized in buffer containing 20 mM HEPES-KOH (HEPES (Fisher BP410); KOH (Sigma 221473), 1 mM EDTA (Sigma E9884), 200 mM sucrose (Sigma S0389), and 1X protease inhibitor (Roche #11873580001). Resulting homogenate was transferred to a 15 mL tube and spun at 600xg for 10-minutes at 4°C to pellet the nucleus. Supernatant was transferred to a new tube and spun at 7,000xg for 10-minutes at 4°C. Supernatant was aspirated and pellet was resuspended in buffer. Spin and pellet washes were repeated for a total of 3X. Final pellet was weighed and promptly stored at - 80°C until further analysis. Mitochondrial enrichment confirmed via Western blot (**Supplemental Figure 3B**).

### Mitochondrial NAD(P)(H) quantification

Pre-weighed mitochondrial pellets were homogenized in volume of 80% MeOH such that final concentration was 50 mg/mL and then diluted 1:50 in 1:1:1:1 PBS:0.2 M NaOH w/ 1% DTAB:0.4M HCl:0.5M Tris (PBS (Gibco 14190250), NaOH (Sigma 221465), DTAB (Sigma D8638), HCl (Avantor 9535) Tris base (Sigma T6066)). NAD(P)(H) was then measured with the Promega NAD/NADH-Glo^TM^ and NADP/NADPH-Glo^TM^ Assays (G9071 and G9081) according to manufacturer instructions.

### Western blots

Protein samples were quantified with Pierce BCA assay (Thermo Scientific 23225). They were then diluted in RIPA buffer (Tris-HCl (Tris base (Sigma T6066), HCl (Avantor 9535)); Triton x-100 (Sigma X100); Na-Deoxycholate (Sigma D6750); SDS (Sigma L3771); NaCl (Sigma, #S7653); EDTA, NaF (Sigma S7920); protease inhibitor (Roche #11873580001)) and sample buffer ((Tris base; SDS; glycerol (Fisher BP229-1); beta-mercaptoethanol; bromophenol blue (Sigma B5525)) and boiled at 95°C for 5-minutes. Samples were run on either 4-20% polyacrylamide gels (Bio-Rad 4561096) or 4-12% Bis-Tris gels (Invitrogen NW04127BOX) at 80 V for 2-hours in either running buffer (Tris base; glycine (RPI G36050); SDS) or buffer (Tris-MOPS-SDS (GenScript M00138)), respectively. Gels were then transferred to PVDF membranes (Bio-Rad #1620177) at 100V for 1.5-hours in transfer buffer (Tris base; glycine; MeOH). Membranes were stained with Ponceau stain (Ponceau S (Sigma P3504), acetic acid (Fisher A38)) for total protein normalization and then blocked with 5% milk (Carnation) in TBST (TBS (Fisher BP2471-1) and Tween-20 (Thermo Scientific J20605.AP)) for 1-hour. After blocking, membranes were washed 4x5-minutes in TBST and incubated in respective primary antibody diluted in 5% BSA (Sigma A9647) overnight at 4°C. The next day, primary antibody was removed, blot was washed 4x5-minutes in TBST, respective secondary antibody was added for 1-hour, and then blot was washed again for 4x5-minutes in TBST. ECL substrate (Bio-Rad #1705062) was then added to the blots and imaged on a Chemi-Doc imager.

### Steroid quantification

Blood samples were collected in EDTA-coated tubes (Fisher NC9990563) and spun at 2,000xg for 15-minutes at 4°C. Resulting plasma was aliquoted into new tubes. Where described, CORT was quantified using Arbor Assay’s Corticosterone Multi-Format ELISA Kit (K014-H1) according to manufacturer’s instructions. Plasma samples were diluted 1:200 in the kit’s assay buffer prior to the start of the assay. For ACTH measurements, plasma samples were diluted 1:3 and quantified using Invitrogen’s Mouse ACTH ELISA Kit (EEL085) according to manufacturer’s instructions.

For the plasma steroid panel, a 25 μL aliquot of serum was added to a 2 mL microcentrifuge tube with 375 μL of water and 20 μL of stable isotope labeled steroid cocktail. The mixture was vortexed for 5-seconds, 200 μL of ZnSO_4_ was added, and then vortexed again for 10-seconds. A 400 μL aliquot of cold methanol was added, samples were vortexed for 30-seconds, then let stand for 5-minutes and then centrifuged at 15,000xg for 10-minutes. Steroids were isolated from the sample using solid phase extraction (Evolute Express ABN (30 mg/1 mL), Biotage, Charlotte, NC). Columns were sequentially conditioned with 1 mL of acetonitrile, 1 mL methanol, and 0.5 mL water. All SPE work was carried out with a positive pressure manifold (Biotage, Charlotte, NC) operated at 6 psi. Samples were loaded onto the conditioned columns and washed with 0.5 mL water and then 0.5 mL of 30% methanol, both of which were discarded. Steroids were eluted with two 0.5 mL aliquots of acetonitrile into a 2 mL polypropylene plate (2 mL Square 96-Well Plate with 100 μL Tapered Reservoir, Analytical Sales, Flanders, NJ). Samples were evaporated under a gentle stream of nitrogen at 40 °C and reconstituted in 50 μL of 50% methanol, vortexed for 30-seconds. Steroids were quantitatively measured using high performance liquid chromatography-mass spectrometry as described ^90,91^.

### Fasting glucose measurements

Fasts began with both the removal of food and cage change to ensure no food was on the bottom of the cage. At the respective timepoint, blood was collected from a tail snip. Glucose was measured with CONTOUR NEXT Blood Glucose Test Strips and CONTOUR NEXT EZ Blood Glucose Monitor.

### Tolerance tests

Mice were fasted for 16-hours from ZT22-ZT14 and time=0 blood glucose was measured as described above. For the glucose tolerance and glucose stimulated insulin secretion tests, mice were given an intraperitoneal (IP) injection of 1.5 g/kg glucose (D-(+)-glucose Sigma G8270). For the insulin tolerance tests, mice were given an IP injection of 0.4 IU/kg. Blood glucose was measured at the indicated timepoints. Insulin was measured from plasma via Crystal Chem’s Ultra Sensitive Mouse Insulin ELISA Kit (#90080).

### Organic and amino acid quantification

Mice were fasted for 16-hours from ZT22-ZT14 or from ZT10-ZT2. At either ZT2 or ZT14, mice were briefly sedated via open-drop method exposure to isoflurane and then sacrificed via decapitation. Livers were immediately excised and flash frozen in liquid nitrogen.

Quantitation of organic acids and amino acids was performed by GC–MS as previously described^57^. Frozen liver samples were immediately spiked with stable isotope-labeled amino acid and organic acid internal standards. Approximately 20 mg of frozen tissue was homogenized in 1 mL of 40:40:20 acetonitrile:methanol containing 0.5% formic acid, vortexed, and incubated on ice for 10 min. Samples were then neutralized by adding 75 μL of 15% NH₄HCO₃, followed by vortexing and an additional 10-minute incubation on ice. After centrifugation at 14,000 RPM for 25-minutes at 4 °C, the supernatant was collected and dried under nitrogen. Next, dried extracts were resuspended in 50 μL of 1% methoxyamine hydrochloride in pyridine and incubated at 37 °C for 90-minutes. Subsequently, 80 μL of MTBSTFA was added, and samples were incubated at 60 °C for 60-minutes. Derivatized samples were transferred to GC vials containing glass inserts and analyzed by GC–MS in selected ion monitoring (SIM) mode. Metabolites were quantified by comparing analyte ion peak areas to those of the corresponding internal standards.

GC-MS analysis was performed using an Agilent 7890-A GC-MS system equipped with an HP-5ms column (30 m × 0.25 mm I.D., 0.25 μm film thickness; Agilent J&W) combined with an Agilent 5975-C mass spectrometer (70eV, electron ionization source). A 1 μL injection volume for all samples was used, and the split mode was adjusted for optimal signal-to-noise ratio.

### De novo lipogenesis

At ZT12, *ad libitum* fed mice were IP injected with 100% ^2^H_2_O (Cambridge Isotope Laboratories #DLM-4-1000) with 0.9% NaCl (Sigma, #S7653) at a dose of 27 μL/g of body weight and then provided with 6% ^2^H_2_O drinking water. At ZT14, mice were briefly sedated via open-drop method exposure to isoflurane and then sacrificed via decapitation. Whole livers were promptly weighed and flash frozen in liquid nitrogen.

Liver DNL was analyzed by HR-Orbitrap-GC/MS as previously described^65,66^. Body water enrichment was measured and used to calculate fractional DNL^65^. Briefly, 20 mg of frozen liver was homogenized in 2 mL tubes with 2.8 mm ceramic beads (Sigma, OMNI-19-628) and 1 mL of MeOH:DCM (1:2, v/v). Tubes were washed twice with 1 mL of MeOH:DCM (1:2), all washes were combined into a new glass tube, and homogenate was spun at 1635×*g* for 5-minutes. Volume equivalent of 1 mg of liver was aliquoted to a new glass tube with 100 μL 0.05 µg/µL ^13^C_16_ palmitate and dried under N_2_ gas. Dried samples were saponified with 2 mL of 0.5 M KOH/MeOH at 80°C for 1-hour. Once the samples cooled, lipids were extracted with 2 mL of DCM and 2 mL of H_2_O and spun at 1635×*g* for 5-minutes. The resulting organic phase was moved to a new glass tube and dried under N_2_ gas. Dried lipid extracts were resuspended in 50 µL of 1% triethylamine/acetone and reacted with 50 µL of 1% PFBBr/acetone for 30-minutes at room temperature. To this solution, 1 mL of isooctane was added before MS analysis.

### Mouse line generation

C57BL/6J background mice harboring the M65A point mutation were generated using CRISPR/Cas9 reagents at the Transgenic Technology Center of UT Southwestern Medical Center. The guide RNAs and donor ssODN were designed to mutate the second in-frame methionine (ATG) in the *Nocturnin* sequence to alanine (GCG). The sgRNA sequence is CCATGGGAAACGGCACCAGTCG. crRNA and tracRNA were annealed and mixed with Cas9 protein to form a ribonucleotide protein complex. The ssODN (Sigma) was added to the mix and the cocktail was microinjected into the cytoplasm of fertilized one-cell eggs isolated from superovulated females. The eggs were incubated in media containing cytochalasin-B immediately before and during microinjection to improve egg survival. Alternatively, CRISPR reagents were delivered to the cytoplasm via electroporation using either a Nepa21 Super Electroporator (NEPAGENE, Ichikawa, Japan) or Gene Pulser (BioRad, Hercules, CA, USA). The surviving eggs were transferred into the oviducts of day 0.5 pseudopregnant recipient ICR females (Envigo, Inc.) to produce putative founder mice. Founder mice were identified via PCR using the primer set 5’-AAATTACCTGCCCGTGTAAGA-3’ and 5’- TGCATTCCTCCAGAAGTTCC -3’ and the amplicon was submitted for Sanger sequencing. F0 mice were bred with C57Bl/6N or J mice to obtain F1 mice heterozygous for the mutated allele. These mice were intercrossed to produce mice homozygous for the M65A mutation.

### Metabolomics analysis

Normalized areas from the metabolomics were used as input for weighted metabolite correlation network analysis using the WGCNA R package^67^. 228 metabolites were used to build a signed network with the blockwiseModules function using the correlation biweight midcorrelation to avoid heavy outliers. The minimum module size was set to 10 and the merging cut height was set to 0.25. A scale free topology was achieved by picking a power of 12 yielding a soft R^2^ of 0.87. Module colors were assigned using labels2colors.

## Acknowledgments

We thank the UTSW Transgenic Technology Core for creating the *Noct^mito^*mice. We thank the UTSW Children’s Research Institute for the circadian NAD(P)(H) and metabolomics measurements. We thank Duyen Pham for her help with the amino acid measurements.

---

**Supplemental Figure 1:**
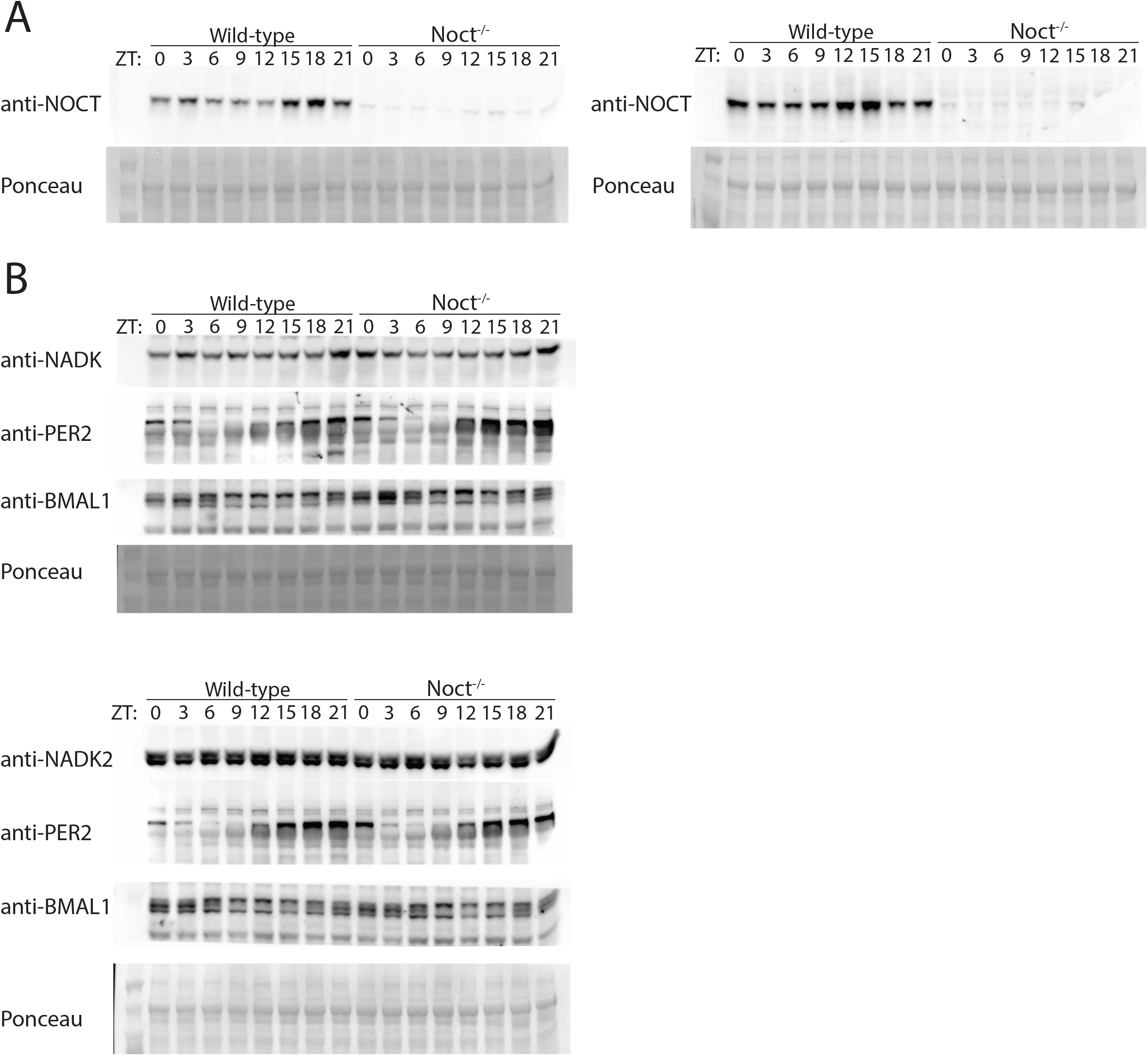
Levels of core clock proteins, NOCT, NADK, and NADK2 in WT and *Noct^-/-^* livers. Western blots of protein extracted from WT and *Noct^-/-^* liver whole cell lysate during the circadian collection outlined in Figure 1A. (A) anti-NOCT and Ponceau stain loading control. (B) anti-NADK, anti-NADK2, anti-PER2, anti-BMAL1, and respective Ponceau stain loading controls.

**Supplemental Figure 2:**
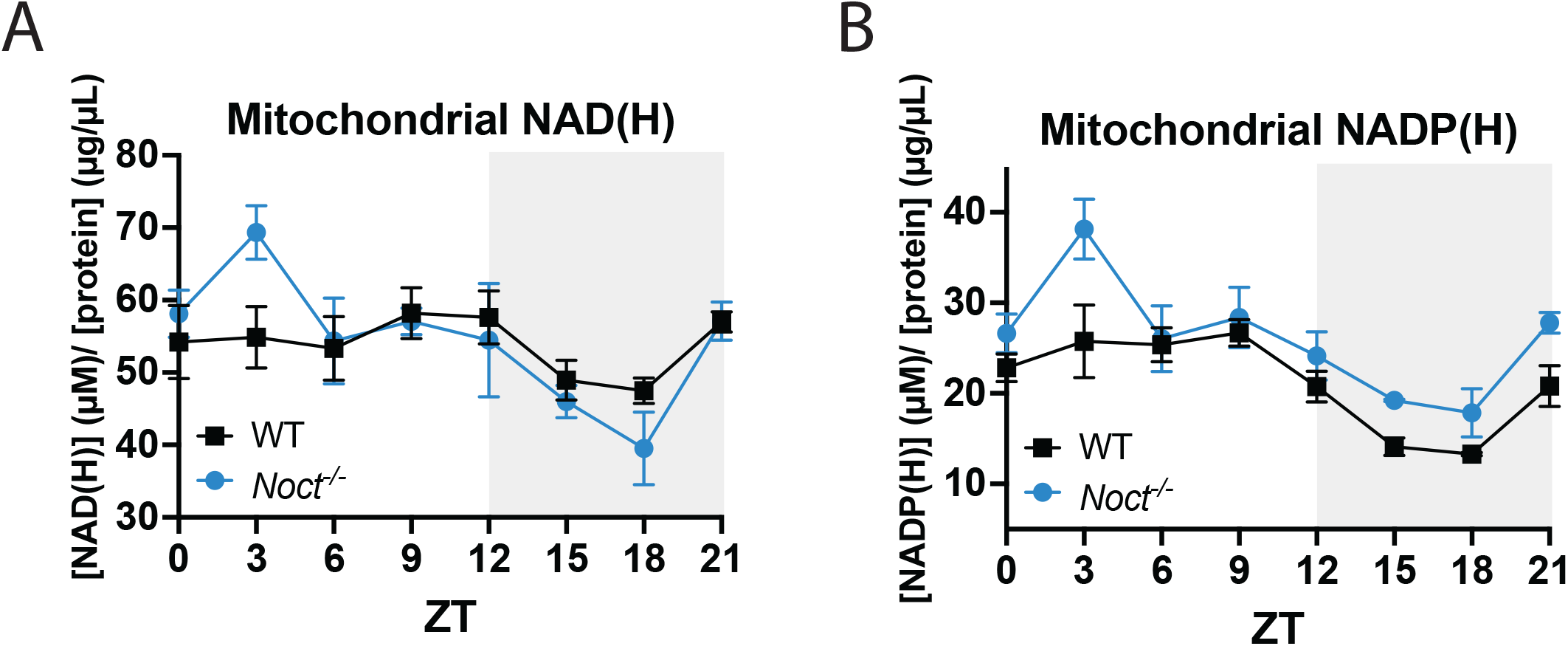
Mitochondrial NAD(H) and NADP(H) in WT and *Noct^-/-^*livers. (A) Mitochondrial NAD(H) concentration normalized to protein concentration. (B) Mitochondrial NADP(H) concentration normalized to protein concentration.

**Supplemental Figure 3:**
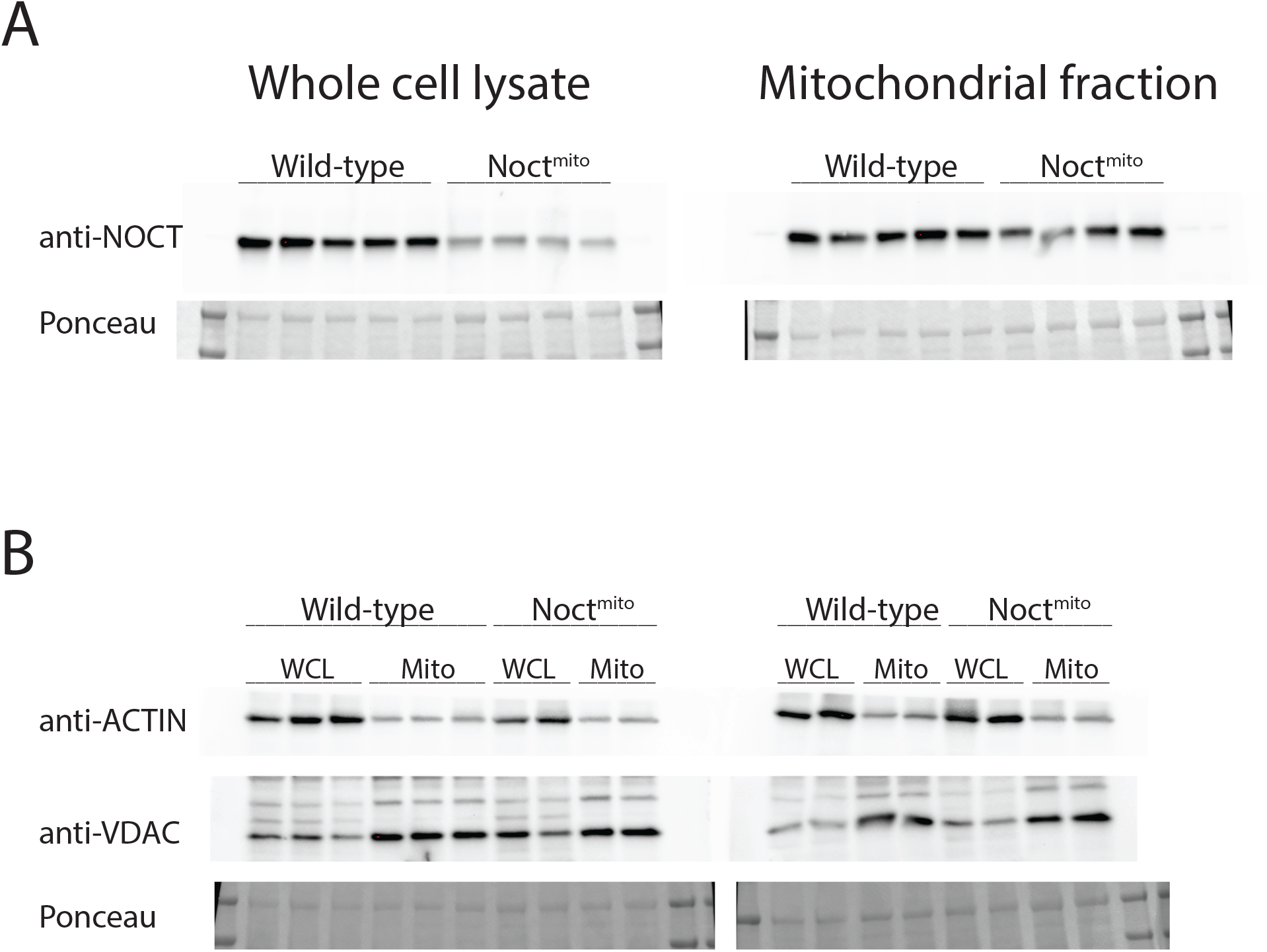
Validation of the *Noct^mito^* mouse line. (A) Western blots of whole cell lysate and isolated mitochondrial fraction from WT and *Noct^mito^*livers probed with anti-NOCT antibody. Ponceau stain of blot as loading control. (B) Western blots of whole cell lysate and isolated mitochondrial fraction from WT and *Noct^mito^* livers probed with anti-ACTIN to show cytosolic depletion in mitochondrial fraction and anti-VDAC to show enrichment on mitochondria in mitochondrial fraction. Ponceau stain of blot as loading control.

**Supplemental Figure 4:**
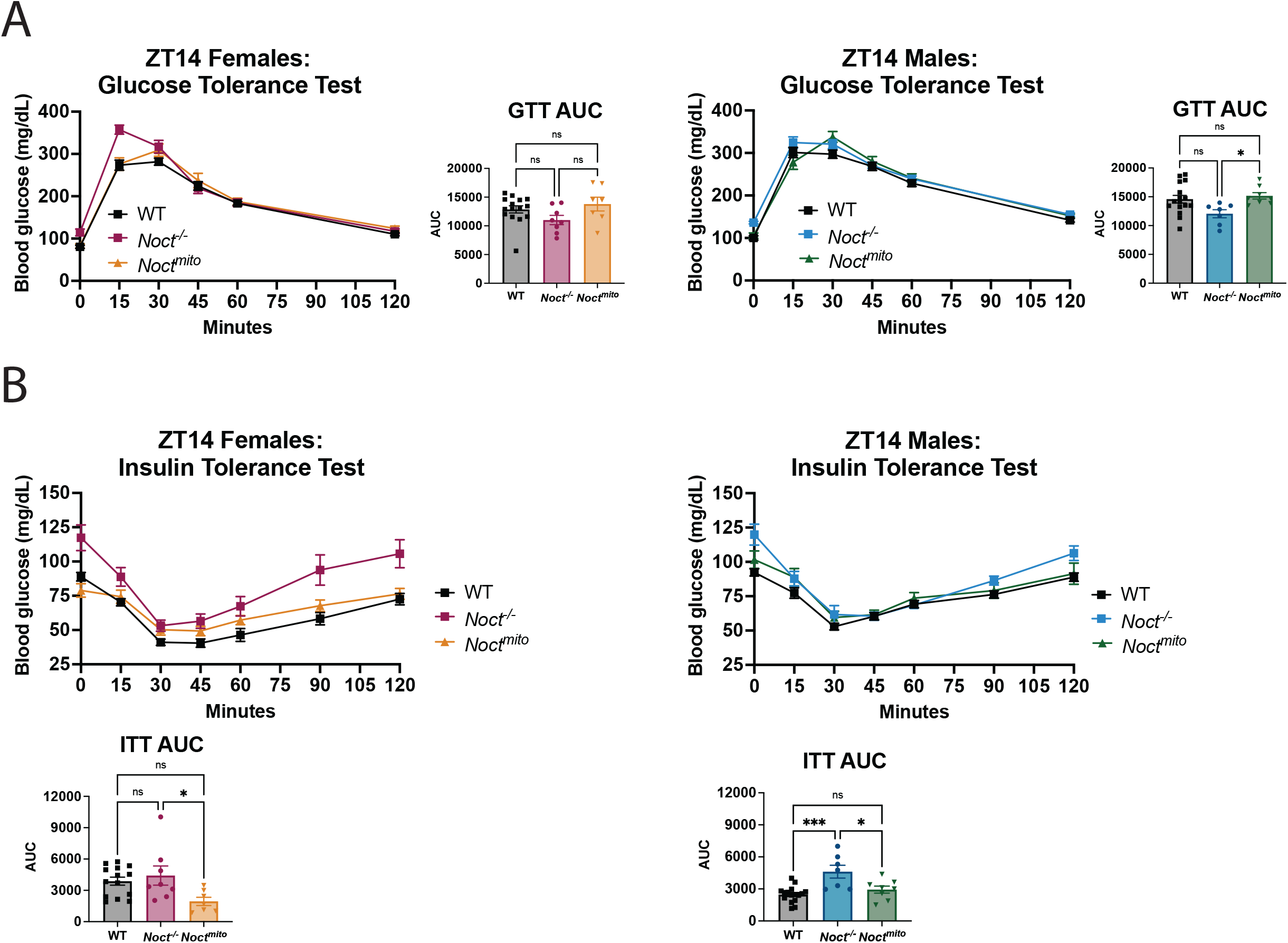
Metabolic phenotyping of the *Noct^mito^* mouse line. (A) IPGTT of male and female WT, *Noct^-/-^*, and *Noct^mito^* mice at ZT14 post 16-hour fast (n=16 for WT, n=7 *Noct^-/-^*, n=8 *Noct^mito^* males; n=15 for WT, n=8 *Noct^-/-^*, n=7 *Noct^mito^* females). Welch’s t-test performed on the AUC. (B) IPITT of male and female WT, *Noct^-/-^*, and *Noct^mito^* mice at ZT14 post 16-hour fast (n=16 for WT, n=7 *Noct^-/-^*, n=8 *Noct^mito^* males; n=14 for WT, n=8 *Noct^-/-^*, n=7 *Noct^mito^* females). Welch’s t-test performed on the AUC. P-value=<0.05 (*), <0.01 (**), <0.001 (***), <0.0001 (****).

**Supplemental Figure 5:**
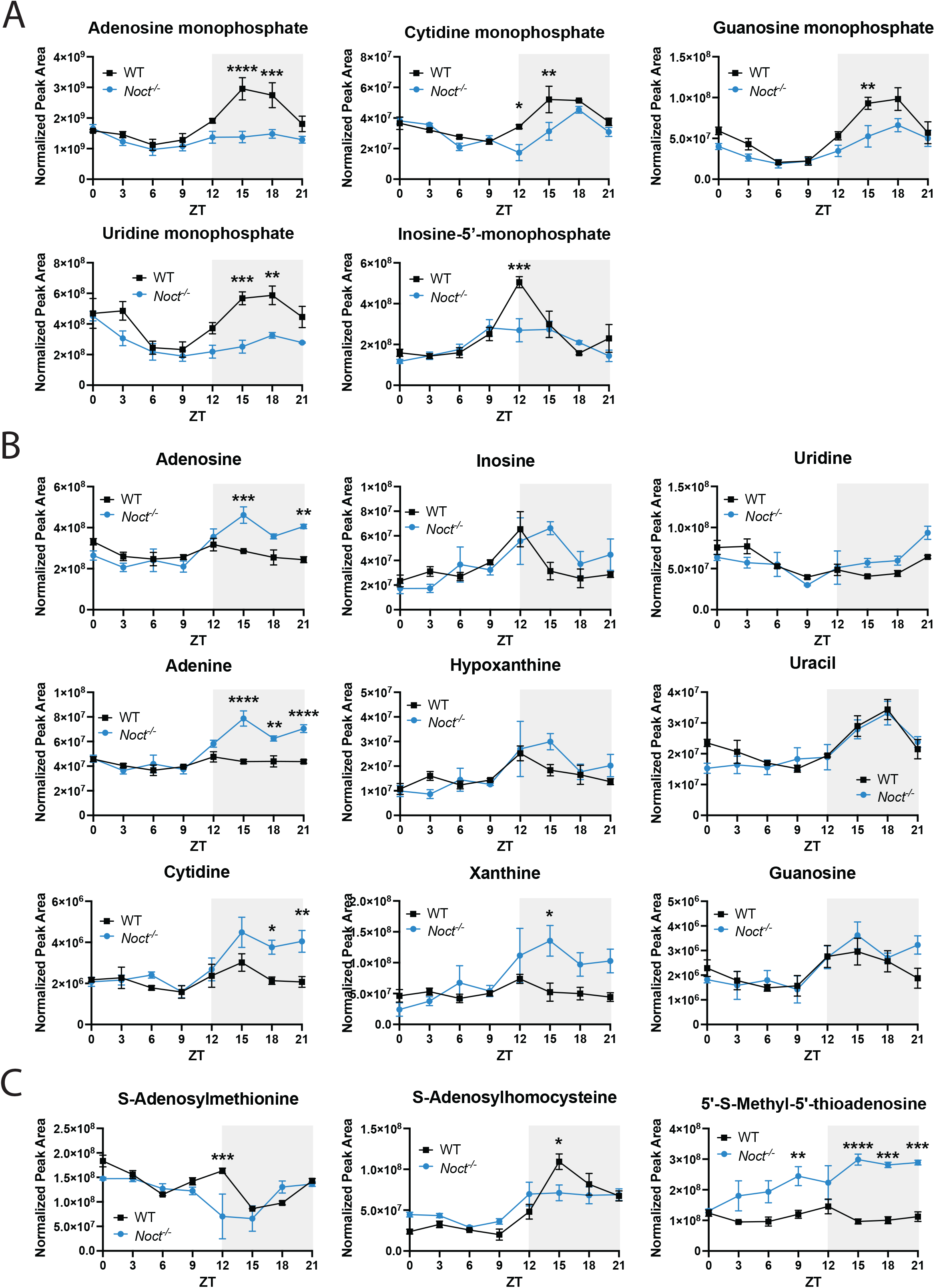
Select metabolites from the circadian untargeted metabolomics dataset. (A) Monophosphates (B) Nucleosides and nucleotides (C) Select one-carbon metabolism metabolites. Two-way ANOVA with multiple comparisons performed on graphs in (A)-(C). P-value=<0.05 (*), <0.01 (**), <0.001 (***), <0.0001 (****).

## Citations

1. Acosta-Rodríguez, V., Rijo-Ferreira, F., Izumo, M., Xu, P., Wight-Carter, M., Green, C.B., and Takahashi, J.S. (2022). Circadian alignment of early onset caloric restriction promotes longevity in male C57BL/6J mice. Science 376, 1192–1202. 10.1126/science.abk0297.

2. Hatori, M., Vollmers, C., Zarrinpar, A., DiTacchio, L., Bushong, Eric A., Gill, S., Leblanc, M., Chaix, A., Joens, M., Fitzpatrick, James A.J., et al. (2012). Time-Restricted Feeding without Reducing Caloric Intake Prevents Metabolic Diseases in Mice Fed a High-Fat Diet. Cell Metabolism 15, 848–860. 10.1016/j.cmet.2012.04.019.

3. Iiams, S.E., Skinner, N.J., Wight-Carter, M., Acosta-Rodríguez, V.A., Green, C.B., and Takahashi, J.S. (2026). Time-restricted feeding extends healthspan in both sexes and lifespan in male C57BL/6 J mice. Nature Aging. 10.1038/s43587-026-01129-8.

4. Green, C.B., Takahashi, J.S., and Bass, J. (2008). The Meter of Metabolism. Cell 134, 728–742. 10.1016/j.cell.2008.08.022.

5. Lévi, F., Zidani, R., and Misset, J.-L. (1997). Randomised multicentre trial of chronotherapy with oxaliplatin, fluorouracil, and folinic acid in metastatic colorectal cancer. The Lancet 350, 681–686. 10.1016/S0140-6736(97)03358-8.

6. Long, J.E., Drayson, M.T., Taylor, A.E., Toellner, K.M., Lord, J.M., and Phillips, A.C. (2016). Morning vaccination enhances antibody response over afternoon vaccination: A cluster-randomised trial. Vaccine 34, 2679–2685. 10.1016/j.vaccine.2016.04.032.

7. Walker, W.H., Bumgarner, J.R., Walton, J.C., Liu, J.A., Meléndez-Fernández, O.H., Nelson, R.J., and DeVries, A.C. (2020). Light Pollution and Cancer. International Journal of Molecular Sciences 21, 9360.

8. Sookoian, S., Gemma, C., Fernández Gianotti, T., Burgueño, A., Alvarez, A., González, C.D., and Pirola, C.J. (2007). Effects of rotating shift work on biomarkers of metabolic syndrome and inflammation. Journal of Internal Medicine 261, 285–292. 10.1111/j.1365-2796.2007.01766.x.

9. Kanki, M., Nath, A.P., Xiang, R., Yiallourou, S., Fuller, P.J., Cole, T.J., Cánovas, R., and Young, M.J. (2023). Poor sleep and shift work associate with increased blood pressure and inflammation in UK Biobank participants. Nature Communications 14, 7096. 10.1038/s41467-023-42758-6.

10. Weed, L., and Zeitzer, J.M. (2025). Circadian-informed modeling predicts regional variation in obesity and stroke outcomes under different permanent US time policies. Proceedings of the National Academy of Sciences 122, e2508293122. doi:10.1073/pnas.2508293122.

11. Parsons, M.J., Moffitt, T.E., Gregory, A.M., Goldman-Mellor, S., Nolan, P.M., Poulton, R., and Caspi, A. (2015). Social jetlag, obesity and metabolic disorder: investigation in a cohort study. Int J Obes (Lond) 39, 842–848. 10.1038/ijo.2014.201.

12. Bass, J., and Takahashi, J.S. (2010). Circadian integration of metabolism and energetics. Science 330, 1349–1354. 10.1126/science.1195027.

13. Sahar, S., and Sassone-Corsi, P. (2012). Regulation of metabolism: the circadian clock dictates the time. Trends Endocrinol Metab 23, 1–8. 10.1016/j.tem.2011.10.005.

14. Chaix, A., Lin, T., Le, H.D., Chang, M.W., and Panda, S. (2019). Time-Restricted Feeding Prevents Obesity and Metabolic Syndrome in Mice Lacking a Circadian Clock. Cell Metabolism 29, 303–319.e304. 10.1016/j.cmet.2018.08.004.

15. Weger, B.D., Gobet, C., David, F.P.A., Atger, F., Martin, E., Phillips, N.E., Charpagne, A., Weger, M., Naef, F., and Gachon, F. (2021). Systematic analysis of differential rhythmic liver gene expression mediated by the circadian clock and feeding rhythms. Proceedings of the National Academy of Sciences 118, e2015803118. doi:10.1073/pnas.2015803118.

16. Goodman, R.P., Calvo, S.E., and Mootha, V.K. (2018). Spatiotemporal compartmentalization of hepatic NADH and NADPH metabolism. Journal of Biological Chemistry 293, 7508–7516. 10.1074/jbc.TM117.000258.

17. Palluth, L., Takahashi, J.S., and Green, C.B. (2025). Keeping up with the nicotinamides: NADP(H), the forgotten circadian cofactor that keeps metabolic time. Life Metabolism. 10.1093/lifemeta/loaf034.

18. Ramsey, K.M., Yoshino, J., Brace, C.S., Abrassart, D., Kobayashi, Y., Marcheva, B., Hong, H.-K., Chong, J.L., Buhr, E.D., Lee, C., et al. (2009). Circadian Clock Feedback Cycle Through NAMPT-Mediated NAD+ Biosynthesis. Science 324, 651–654. doi:10.1126/science.1171641.

19. Peek, C.B., Affinati, A.H., Ramsey, K.M., Kuo, H.Y., Yu, W., Sena, L.A., Ilkayeva, O., Marcheva, B., Kobayashi, Y., Omura, C., et al. (2013). Circadian clock NAD+ cycle drives mitochondrial oxidative metabolism in mice. Science 342, 1243417. 10.1126/science.1243417.

20. Hepler, C., Waldeck, N.J., Weidemann, B.J., Marcheva, B., Chen, Y.-J., Hecker, J., Zhu, Z., Nozawa, R., Mastroni, J.V., Thorne, A.K., et al. (2026). Adipocyte NADH dehydrogenase reverses circadian and diet-induced metabolic syndrome. Nature Metabolism 8, 559–571. 10.1038/s42255-026-01464-5.

21. Levine, D.C., Kuo, H.Y., Hong, H.K., Cedernaes, J., Hepler, C., Wright, A.G., Sommars, M.A., Kobayashi, Y., Marcheva, B., Gao, P., et al. (2021). NADH inhibition of SIRT1 links energy state to transcription during time-restricted feeding. Nat Metab 3, 1621–1632. 10.1038/s42255-021-00498-1.

22. Estrella, M.A., Du, J., Chen, L., Rath, S., Prangley, E., Chitrakar, A., Aoki, T., Schedl, P., Rabinowitz, J., and Korennykh, A. (2019). The metabolites NADP+ and NADPH are the targets of the circadian protein Nocturnin (Curled). Nature Communications 10, 2367. 10.1038/s41467-019-10125-z.

23. Olivas-Rasmussen, C., Palluth, L., Rasmussen, E.S., and Green, C.B. (2026). Characterization of the enzymatic activity of the rhythmic human NADP(H) phosphatase, Nocturnin. Archives of Biochemistry and Biophysics 776, 110692. 10.1016/j.abb.2025.110692.

24. Green, C.B., and Besharse, J.C. (1996). Identification of a novel vertebrate circadian clock-regulated gene encoding the protein nocturnin. Proc Natl Acad Sci U S A 93, 14884–14888. 10.1073/pnas.93.25.14884.

25. Wang, Y., Osterbur, D.L., Megaw, P.L., Tosini, G., Fukuhara, C., Green, C.B., and Besharse, J.C. (2001). Rhythmic expression of Nocturnin mRNA in multiple tissues of the mouse. BMC Dev Biol 1, 9. 10.1186/1471-213x-1-9.

26. Green, C.B., and Besharse, J.C. (1996). Use of a high stringency differential display screen for identification of retinal mRNAs that are regulated by a circadian clock. Molecular Brain Research 37, 157–165. 10.1016/0169-328X(95)00307-E.

27. Green, C.B., Douris, N., Kojima, S., Strayer, C.A., Fogerty, J., Lourim, D., Keller, S.R., and Besharse, J.C. (2007). Loss of Nocturnin, a circadian deadenylase, confers resistance to hepatic steatosis and diet-induced obesity. Proceedings of the National Academy of Sciences 104, 9888–9893. 10.1073/pnas.0702448104.

28. Douris, N., Kojima, S., Pan, X., Lerch-Gaggl, A.F., Duong, S.Q., Hussain, M.M., and Green, C.B. (2011). Nocturnin regulates circadian trafficking of dietary lipid in intestinal enterocytes. Curr Biol 21, 1347–1355. 10.1016/j.cub.2011.07.018.

29. Stubblefield, J.J., Gao, P., Kilaru, G., Mukadam, B., Terrien, J., and Green, C.B. (2018). Temporal Control of Metabolic Amplitude by Nocturnin. Cell Rep 22, 1225–1235. 10.1016/j.celrep.2018.01.011.

30. Wickramaratne, A.C., Li, L., Hopkins, J.B., Joachimiak, L.A., and Green, C.B. (2022). The Disordered Amino Terminus of the Circadian Enzyme Nocturnin Modulates Its NADP(H) Phosphatase Activity by Changing Protein Dynamics. Biochemistry 61, 1091–1102. 10.1021/acs.biochem.2c00072.

31. Laothamatas, I., Gao, P., Wickramaratne, A., Quintanilla, C.G., Dino, A., Khan, C.A., Liou, J., and Green, C.B. (2020). Spatiotemporal regulation of NADP(H) phosphatase Nocturnin and its role in oxidative stress response. Proc Natl Acad Sci U S A 117, 993–999. 10.1073/pnas.1913712117.

32. Onder, Y., Laothamatas, I., Berto, S., Sewart, K., Kilaru, G., Bordieanu, B., Stubblefield, J.J., Konopka, G., Mishra, P., and Green, C.B. (2019). The Circadian Protein Nocturnin Regulates Metabolic Adaptation in Brown Adipose Tissue. iScience 19, 83–92. 10.1016/j.isci.2019.07.016.

33. Lu, W., Wang, L., Chen, L., Hui, S., and Rabinowitz, J.D. (2018). Extraction and Quantitation of Nicotinamide Adenine Dinucleotide Redox Cofactors. Antioxid Redox Signal 28, 167–179. 10.1089/ars.2017.7014.

34. Aurora, A.B., Khivansara, V., Leach, A., Gill, J.G., Martin-Sandoval, M., Yang, C., Kasitinon, S.Y., Bezwada, D., Tasdogan, A., Gu, W., et al. (2022). Loss of glucose 6-phosphate dehydrogenase function increases oxidative stress and glutaminolysis in metastasizing melanoma cells. Proc Natl Acad Sci U S A 119. 10.1073/pnas.2120617119.

35. O’Neill, J.S., and Reddy, A.B. (2011). Circadian clocks in human red blood cells. Nature 469, 498–503. 10.1038/nature09702.

36. Luongo, T.S., Eller, J.M., Lu, M.-J., Niere, M., Raith, F., Perry, C., Bornstein, M.R., Oliphint, P., Wang, L., McReynolds, M.R., et al. (2020). SLC25A51 is a mammalian mitochondrial NAD+ transporter. Nature 588, 174–179. 10.1038/s41586-020-2741-7.

37. Ryu, K.W., Nandu, T., Kim, J., Challa, S., DeBerardinis, R.J., and Kraus, W.L. (2018). Metabolic regulation of transcription through compartmentalized NAD^+^ biosynthesis. Science 360, eaan5780. doi:10.1126/science.aan5780.

38. Cambronne, X.A., Stewart, M.L., Kim, D., Jones-Brunette, A.M., Morgan, R.K., Farrens, D.L., Cohen, M.S., and Goodman, R.H. (2016). Biosensor reveals multiple sources for mitochondrial NAD^+^. Science 352, 1474–1477. doi:10.1126/science.aad5168.

39. Cambronne, X.A., and Kraus, W.L. (2020). Location, Location, Location: Compartmentalization of NAD+ Synthesis and Functions in Mammalian Cells. Trends in Biochemical Sciences 45, 858–873. 10.1016/j.tibs.2020.05.010.

40. Sjöberg, B., de la Torre, B., Hedman, M., Falkay, G., and Diczfalusy, E. (1979). Circadian variation in systemic hormone levels in healthy men. Journal of Endocrinological Investigation 2, 131–137. 10.1007/BF03349304.

41. de Peretti, E., and Mappus, E. (1983). Pattern of plasma pregnenolone sulfate levels in humans from birth to adulthood. J Clin Endocrinol Metab 57, 550–556. 10.1210/jcem-57-3-550.

42. Le Minh, N., Damiola, F., Tronche, F., Schütz, G., and Schibler, U. (2001). Glucocorticoid hormones inhibit food-induced phase-shifting of peripheral circadian oscillators. The EMBO Journal 20, 7128–7136. 10.1093/emboj/20.24.7128.

43. Reddy, A.B., Maywood, E.S., Karp, N.A., King, V.M., Inoue, Y., Gonzalez, F.J., Lilley, K.S., Kyriacou, C.P., and Hastings, M.H. (2007). Glucocorticoid signaling synchronizes the liver circadian transcriptome. Hepatology 45, 1478–1488. 10.1002/hep.21571.

44. Oster, H., Damerow, S., Kiessling, S., Jakubcakova, V., Abraham, D., Tian, J., Hoffmann, M.W., and Eichele, G. (2006). The circadian rhythm of glucocorticoids is regulated by a gating mechanism residing in the adrenal cortical clock. Cell Metabolism 4, 163–173. 10.1016/j.cmet.2006.07.002.

45. De Boer, S.F., and Van Der Gugten, J. (1987). Daily variations in plasma noradrenaline, adrenaline and corticosterone concentrations in rats. Physiology & Behavior 40, 323–328. 10.1016/0031-9384(87)90054-0.

46. Balsalobre, A., Brown, S.A., Marcacci, L., Tronche, F., Kellendonk, C., Reichardt, H.M., Schütz, G., and Schibler, U. (2000). Resetting of Circadian Time in Peripheral Tissues by Glucocorticoid Signaling. Science 289, 2344–2347. doi:10.1126/science.289.5488.2344.

47. Karatsoreos, I.N., Bhagat, S.M., Bowles, N.P., Weil, Z.M., Pfaff, D.W., and McEwen, B.S. (2010). Endocrine and Physiological Changes in Response to Chronic Corticosterone: A Potential Model of the Metabolic Syndrome in Mouse. Endocrinology 151, 2117–2127. 10.1210/en.2009-1436.

48. Peckett, A.J., Wright, D.C., and Riddell, M.C. (2011). The effects of glucocorticoids on adipose tissue lipid metabolism. Metabolism 60, 1500–1510. 10.1016/j.metabol.2011.06.012.

49. Weitzman, E.D., Fukushima, D., Nogeire, C., Roffwarg, H., Gallagher, T.F., and Hellman, L. (1971). Twenty-four Hour Pattern of the Episodic Secretion of Cortisol in Normal Subjects. The Journal of Clinical Endocrinology & Metabolism 33, 14–22. 10.1210/jcem-33-1-14.

50. Kahnt, F.W., Milani, A., Steffen, H., and Neher, R. (1974). The Rate-Limiting Step of Adrenal Steroidogenesis and Adenosine 3′:5′-Monophosphate. European Journal of Biochemistry 44, 243–250. 10.1111/j.1432-1033.1974.tb03479.x.

51. Miller, W.L., and Auchus, R.J. (2011). The Molecular Biology, Biochemistry, and Physiology of Human Steroidogenesis and Its Disorders. Endocrine Reviews 32, 81–151. 10.1210/er.2010-0013.

52. Payne, A.H., and Hales, D.B. (2004). Overview of Steroidogenic Enzymes in the Pathway from Cholesterol to Active Steroid Hormones. Endocrine Reviews 25, 947–970. 10.1210/er.2003-0030.

53. Dunford, E.C., and Riddell, M.C. (2016). The Metabolic Implications of Glucocorticoids in a High-Fat Diet Setting and the Counter-Effects of Exercise. Metabolites 6, 44.

54. Ferraù, F., and Korbonits, M. (2015). Metabolic comorbidities in Cushing’s syndrome. European Journal of Endocrinology 173, M133–M157. 10.1530/eje-15-0354.

55. Salehidoost, R., and Korbonits, M. (2022). Glucose and lipid metabolism abnormalities in Cushing’s syndrome. Journal of Neuroendocrinology 34, e13143. 10.1111/jne.13143.

56. Tannenbaum, B.M., Brindley, D.N., Tannenbaum, G.S., Dallman, M.F., McArthur, M.D., and Meaney, M.J. (1997). High-fat feeding alters both basal and stress-induced hypothalamic-pituitary-adrenal activity in the rat. American Journal of Physiology-Endocrinology and Metabolism 273, E1168–E1177. 10.1152/ajpendo.1997.273.6.E1168.

57. Deja, S., Fletcher, J.A., Kim, C.-W., Kucejova, B., Fu, X., Mizerska, M., Villegas, M., Pudelko-Malik, N., Browder, N., Inigo-Vollmer, M., et al. (2024). Hepatic malonyl-CoA synthesis restrains gluconeogenesis by suppressing fat oxidation, pyruvate carboxylation, and amino acid availability. Cell Metabolism 36, 1088–1104.e1012. 10.1016/j.cmet.2024.02.004.

58. Horiuchi, T., Kaneko, K., Hosaka, S., Uno, K., Tomiyama, S., Takahashi, K., Yamato, M., Endo, A., Sugawara, H., Kawana, Y., et al. (2025). Redox-dependent liver gluconeogenesis impacts different intensity exercise in mice. Nature Metabolism 7, 1991–2003. 10.1038/s42255-025-01373-z.

59. Madiraju, A.K., Erion, D.M., Rahimi, Y., Zhang, X.-M., Braddock, D.T., Albright, R.A., Prigaro, B.J., Wood, J.L., Bhanot, S., MacDonald, M.J., et al. (2014). Metformin suppresses gluconeogenesis by inhibiting mitochondrial glycerophosphate dehydrogenase. Nature 510, 542–546. 10.1038/nature13270.

60. Sugano, T., Shiota, M., Tanaka, T., Miyamae, Y., Shimada, M., and Oshino, N. (1980). Intracellular Redox State and Stimulation of Gluconeogenesis by Glucagon and Norepinephrine in the Perfused Rat Liver. The Journal of Biochemistry 87, 153–166. 10.1093/oxfordjournals.jbchem.a132721.

61. Berdanier, C.D. (1989). Role of glucocorticoids in the regulation of lipogenesis. The FASEB Journal 3, 2179–2183. 10.1096/fasebj.3.10.2666232.

62. Wang, Y., Urs, S., Kim, S., Soltani-Bejnood, M., Quigley, N., Heo, Y.-R., Standridge, M., Andersen, B., Joshi, R., Wortman, P., et al. (2004). The Human Fatty Acid Synthase Gene and De Novo Lipogenesis Are Coordinately Regulated in Human Adipose Tissue. The Journal of Nutrition 134, 1032–1038. 10.1093/jn/134.5.1032.

63. Minshull, M., and Strong, C.R. (1985). The stimulation of lipogenesis in white adipose tissue from fed rats by corticosterone. The International journal of biochemistry 17, 529–532. 10.1016/0020-711x(85)90151-x.

64. Campbell, J.E., Peckett, A.J., D’souza, A.M., Hawke, T.J., and Riddell, M.C. (2011). Adipogenic and lipolytic effects of chronic glucocorticoid exposure. American Journal of Physiology-Cell Physiology 300, C198–C209. 10.1152/ajpcell.00045.2010.

65. Fu, X., Deja, S., Fletcher, J.A., Anderson, N.N., Mizerska, M., Vale, G., Browning, J.D., Horton, J.D., McDonald, J.G., Mitsche, M.A., and Burgess, S.C. (2021). Measurement of lipogenic flux by deuterium resolved mass spectrometry. Nature Communications 12, 3756. 10.1038/s41467-021-23958-4.

66. Rose, J.P., Morgan, D.A., Sullivan, A.I., Fu, X., Inigo-Vollmer, M., Burgess, S.C., Meyerholz, D.K., Rahmouni, K., and Potthoff, M.J. (2025). FGF21 reverses MASH through coordinated actions on the CNS and liver. Cell Metabolism 37, 1515–1529.e1516. 10.1016/j.cmet.2025.04.014.

67. Langfelder, P., and Horvath, S. (2008). WGCNA: an R package for weighted correlation network analysis. BMC Bioinformatics 9, 559. 10.1186/1471-2105-9-559.

68. Sebo, Z.L., Chakrabarty, R.P., Grant, R.A., D’Alessandro, K.B., Koss, A.R., Blum, J.L.E., Davidson, S.M., Reczek, C.R., and Chandel, N.S. (2026). Metformin inhibits mitochondrial complex I in intestinal epithelium to promote glycaemic control. Nature Metabolism. 10.1038/s42255-026-01530-y.

69. Shen, K., Pender, C.L., Bar-Ziv, R., Zhang, H., Wickham, K., Willey, E., Durieux, J., Ahmad, Q., and Dillin, A. (2022). Mitochondria as Cellular and Organismal Signaling Hubs. Annu Rev Cell Dev Biol 38, 179–218. 10.1146/annurev-cellbio-120420-015303.

70. Meichsner, A., Bader, V., and Winklhofer, K.F. (2026). Mitochondria as sources and targets of cellular signaling. Molecular Cell 86, 503–521. 10.1016/j.molcel.2026.01.008.

71. Kory, N., uit de Bos, J., van der Rijt, S., Jankovic, N., Güra, M., Arp, N., Pena, I.A., Prakash, G., Chan, S.H., Kunchok, T., et al. (2020). MCART1/SLC25A51 is required for mitochondrial NAD transport. Science Advances 6, eabe5310. doi:10.1126/sciadv.abe5310.

72. Davila, A., Liu, L., Chellappa, K., Redpath, P., Nakamaru-Ogiso, E., Paolella, L.M., Zhang, Z., Migaud, M.E., Rabinowitz, J.D., and Baur, J.A. (2018). Nicotinamide adenine dinucleotide is transported into mammalian mitochondria. eLife 7, e33246. 10.7554/eLife.33246.

73. Høyland, L.E., VanLinden, M.R., Niere, M., Strømland, Ø., Sharma, S., Dietze, J., Tolås, I., Lucena, E., Bifulco, E., Sverkeli, L.J., et al. (2024). Subcellular NAD(+) pools are interconnected and buffered by mitochondrial NAD(). Nat Metab 6, 2319–2337. 10.1038/s42255-024-01174-w.

74. Ohashi, K., Kawai, S., Koshimizu, M., and Murata, K. (2011). NADPH regulates human NAD kinase, a NADP⁺-biosynthetic enzyme. Mol Cell Biochem 355, 57–64. 10.1007/s11010-011-0838-x.

75. Pollak, N., Niere, M., and Ziegler, M. (2007). NAD Kinase Levels Control the NADPH Concentration in Human Cells *. Journal of Biological Chemistry 282, 33562–33571. 10.1074/jbc.M704442200.

76. Kim, D., Kesavan, R., Ryu, K., Dey, T., Marckx, A., Menezes, C., Praharaj, P.P., Morley, S., Ko, B., Soflaee, M.H., et al. (2025). Mitochondrial NADPH fuels mitochondrial fatty acid synthesis and lipoylation to power oxidative metabolism. Nature Cell Biology 27, 790–800. 10.1038/s41556-025-01655-4.

77. Yang, R., Yang, C., Ma, L., Zhao, Y., Guo, Z., Niu, J., Chu, Q., Ma, Y., and Li, B. (2022). Identification of purine biosynthesis as an NADH-sensing pathway to mediate energy stress. Nature Communications 13, 7031. 10.1038/s41467-022-34850-0.

78. Singh, C., Jin, B., Shrestha, N., Markhard, A.L., Panda, A., Calvo, S.E., Deik, A., Pan, X., Zuckerman, A.L., Ben Saad, A., et al. (2024). ChREBP is activated by reductive stress and mediates GCKR-associated metabolic traits. Cell Metabolism 36, 144–158.e147. 10.1016/j.cmet.2023.11.010.

79. Luengo, A., Li, Z., Gui, D.Y., Sullivan, L.B., Zagorulya, M., Do, B.T., Ferreira, R., Naamati, A., Ali, A., Lewis, C.A., et al. (2021). Increased demand for NAD+ relative to ATP drives aerobic glycolysis. Molecular Cell 81, 691–707.e696. 10.1016/j.molcel.2020.12.012.

80. Rutter, J., Reick, M., Wu, L.C., and McKnight, S.L. (2001). Regulation of Clock and NPAS2 DNA Binding by the Redox State of NAD Cofactors. Science 293, 510–514. doi:10.1126/science.1060698.

81. Iiams, S.E., Skinner, N.J., Green, C.B., and Takahashi, J.S. (2026). Circadian Alignment Through Time-Restricted Feeding: Implications for Health and Longevity. Annual Review of Nutrition. 10.1146/annurev-nutr-112525-011241.

82. Makris, K., Fonda, V., Ramadhani, F.F., Fadel, L., Davezac, M., Payet, B., Deligiannis, I.K., Zhang, L., Horn, T., Heimerl, L., et al. (2025). Hepatic metabolic reprogramming in male mice during short-term caloric restriction involves enhanced glucocorticoid rhythms. Nature Communications 16, 11106. 10.1038/s41467-025-67228-z.

83. Tholen, S., Patel, R., Agas, A., Kovary, K.M., Rabiee, A., Nicholls, H.T., Bielczyk-Maczyńska, E., Yang, W., Kraemer, F.B., and Teruel, M.N. (2022). Flattening of circadian glucocorticoid oscillations drives acute hyperinsulinemia and adipocyte hypertrophy. Cell Reports 39. 10.1016/j.celrep.2022.111018.

84. Bahrami-Nejad, Z., Zhao, M.L., Tholen, S., Hunerdosse, D., Tkach, K.E., van Schie, S., Chung, M., and Teruel, M.N. (2018). A Transcriptional Circuit Filters Oscillating Circadian Hormonal Inputs to Regulate Fat Cell Differentiation. Cell Metabolism 27, 854–868.e858. 10.1016/j.cmet.2018.03.012.

85. Kroon, J., Schilperoort, M., In het Panhuis, W., van den Berg, R., van Doeselaar, L., Verzijl, C.R.C., van Trigt, N., Mol, I.M., Sips, H.H.C.M., van den Heuvel, J.K., et al. (2021). A physiological glucocorticoid rhythm is an important regulator of brown adipose tissue function. Molecular Metabolism 47, 101179. 10.1016/j.molmet.2021.101179.

86. Schilperoort, M., Kroon, J., Kooijman, S., Smit, A.E., Gentenaar, M., Mletzko, K., Schmidt, F.N., van Ruijven, L., Busse, B., Pereira, A.M., et al. (2021). Loss of glucocorticoid rhythm induces an osteoporotic phenotype in female mice. Aging Cell 20, e13474. 10.1111/acel.13474.

87. Lambertucci, F., Castinetti, F., Martins, I., López-Otín, C., and Kroemer, G. Pro-aging effects of chronic glucocorticoid signaling. Cell Metabolism. 10.1016/j.cmet.2026.05.002.

88. DeVilbiss, A.W., Zhao, Z., Martin-Sandoval, M.S., Ubellacker, J.M., Tasdogan, A., Agathocleous, M., Mathews, T.P., and Morrison, S.J. (2021). Metabolomic profiling of rare cell populations isolated by flow cytometry from tissues. Elife 10. 10.7554/eLife.61980.

89. Tasdogan, A., Faubert, B., Ramesh, V., Ubellacker, J.M., Shen, B., Solmonson, A., Murphy, M.M., Gu, Z., Gu, W., Martin, M., et al. (2020). Metabolic heterogeneity confers differences in melanoma metastatic potential. Nature 577, 115–120. 10.1038/s41586-019-1847-2.

90. Braun, V., Stuppner, H., Risch, L., and Seger, C. (2022). Design and Validation of a Sensitive Multisteroid LC-MS/MS Assay for the Routine Clinical Use: One-Step Sample Preparation with Phospholipid Removal and Comparison to Immunoassays. Int J Mol Sci 23 10.3390/ijms232314691.

91. Peitzsch, M., Dekkers, T., Haase, M., Sweep, F.C., Quack, I., Antoch, G., Siegert, G., Lenders, J.W., Deinum, J., Willenberg, H.S., and Eisenhofer, G. (2015). An LC-MS/MS method for steroid profiling during adrenal venous sampling for investigation of primary aldosteronism. J Steroid Biochem Mol Biol 145, 75–84. 10.1016/j.jsbmb.2014.10.006.

